# 3D-imaging reveals conserved cerebrospinal fluid drainage via meningeal lymphatic vasculature in mice and humans

**DOI:** 10.1101/2022.01.13.476230

**Authors:** Laurent Jacob, Jose de Brito Neto, Stephanie Lenck, Celine Corcy, Farhat Benbelkacem, Luiz Henrique Geraldo, Yunling Xu, Jean-Mickael Thomas, Marie-Renee El Kamouh, Marie-Claude Potier, Stephane Haik, Stephane Lehericy, Anne Eichmann, Jean-Leon Thomas

**Author notes:** contributed equally (Jose de Brito Neto, Stephanie Lenck).

## Abstract

Meningeal lymphatic vessels (MLVs) contribute to waste product elimination and immune surveillance in brain tissues. MLVs were identified in the dorsal and caudo-basal regions of the dura mater, where they ensure the clearance of cerebrospinal fluid (CSF). Whether MLVs exist in the complex anterior part of the murine and human skull, and how they connect with the glymphatic system and extracranial lymphatic vasculature remained unclear. Here, we generated three-dimensional (3D) maps of MLV drainage by light-sheet fluorescence microscopy (LSFM) imaging of mouse whole-head preparations following fluorescent OVA-A^555^ tracer injections into the CSF. In humans, we performed real-time magnetic resonance vessel wall imaging (MR-VWI) after systemic gadobutrol injections. We observed a conserved 3D-anatomy of MLVs in mice and humans, and we discovered an extended anterior network around the dural cavernous sinus including multiple capillary beds and exit routes through the foramina of emissary veins. MR-VWI may provide a diagnostic tool for patients with CSF drainage defects and neurological diseases.

**Short abstract:** We established the 3D-anatomy of meningeal lymphatic vasculature and associated CSF drainage by postmortem light-sheet imaging in mice and by real-time magnetic resonance imaging in humans, demonstrating conserved lymphatic circuitries in contact with dural venous sinuses.

## Introduction

MLVs are located in the dura mater of the brain and spinal cord of various vertebrate species, including humans (Aspelund et al., 2015; Louveau et al., 2015; Absinta et al., 2017; Antila et al., 2017; Jacob et al., 2019). MLVs transport macromolecules and antigens from the cerebrospinal fluid (CSF) and interstitial fluids (ISF) of the central nervous system (CNS) to CNS-draining lymph nodes. MLVs bridge the CNS with the peripheral immune system and thereby control both brain waste clearance and neuro-immune communication. MLVs were (re-)discovered in 2015 (Louveau et al., 2015; Aspelund et al., 2015) and have since been shown to play fundamental roles in various mouse models of CNS pathologies, such as multiple sclerosis, Alzheimer’s disease, ischemic stroke and brain tumors (Louveau et al., 2018; Da Mesquita et al., 2018; Hsu et al., 2019; Esposito et al., 2019; Song et al., 2020; Hu et al., 2020). MLVs have thus become attractive as potential therapeutic targets against CNS diseases (Song et al., 2020; Louveau et al., 2016).

The contribution of MLVs to the drainage of CSF outflow from the skull and the clearance of solute waste from CNS tissues has been debated (Louveau et al., 2015; Da Mesquita et al., 2018). Perineural and perivascular spaces were long considered to be the main pathway of CSF outflow into extracranial lymphatics (Tarasoff-Conway et al., 2015; Engelhardt et al., 2016). CSF outflow was shown to drain within sheaths of cranial nerves to reach extracranial lymphatics and collecting lymph nodes (LNs) in the neck (McComb, 1983; Bradbury and Cserr, 1985; Koh et al., 2005). In addition, elegant imaging approaches using near-infrared or dynamic contrast-enhanced magnetic resonance (Ma et al., 2017, 2019) also showed CSF bulk outflow into extracranial lymphatic pathways that proceeds through the basal cisterns, which are the expansions of the subarachnoid space prolonging around cranial nerves and intracranial vessels (Altafulla et al., 2019). However, an additional contribution of dural lymphatics to CSF drainage was revealed by tracer injections into the CSF, which were taken up by MLVs (Antila et al., 2017; Louveau et al., 2018; Hsu et al., 2019; Ahn et al., 2019). Thus, experimental data support the contribution of MLVs to CSF drainage and warrant a thorough evaluation across the whole dura mater in mice and humans.

The CSF is continuously produced by the choroid plexus, and circulates through the CNS internal ventricles, the subarachnoid space and cisterns, as well as along the perivascular spaces of cerebral arteries and veins (Esposito et al., 2019). CSF exchanges with the ISF of the neuropil through the astroglial glymphatic system, thereby facilitating waste clearance from the CNS (Iliff et al., 2012; Hu et al., 2020; Song et al., 2020; Yanev et al., 2020). The glymphatic system generates an outflow of CNS-derived fluids and waste solutes (CSF/ISF) that will subsequently drain out of the skull and the vertebral canal. In the meninges, the glymphatic outflow exits in the perivascular spaces of cerebral veins which converge into the venous sinuses of the dura mater. Dural venous sinuses are key sites of CNS antigen sampling and immune cell egression from the blood into the meninges (Rustenhoven et al., 2021). Dural sinuses also neighbor the MLVs, and the focal ablation of MLVs impaired the glymphatic clearance of toxic aggregates in mice (Da Mesquita et al., 2021). MLVs are thus ideally located to collect brain clearance products at exit points of the perivenous spaces, dowstream of the glymphatic system. However, lymphatic uptake and CSF/ISF drainage pathways from the dura to the collecting LNs remained to be established.

In fact, MLVs have been less explored for their overall architecture and functional organization than for their physiology and pathophysiology. MLV anatomy has been carefully described in the dura mater of the calvaria (Louveau et al., 2015; Aspelund et al., 2015) and in the posterior fossa of the skull base (Antila et al., 2017; Ahn et al., 2019), identifying lymphatic drainage pathways in the dorsal and caudo-basal parts of the skull. Whether additional outflow tracts existed in other parts of the skull remained poorly documented. For example, in the anterior part of the dural sinus system, the cavernous sinus has been suggested to participate in CSF absorption (Koh et al., 2005). The cavernous sinus collects blood from the superficial and deep middle cerebral veins and the ophtalmic and facial regions (Haines, 2018). CSF drainage from this anterior region has been identified (Antila et al., 2017; Ma et al., 2017; Decker et al., 2021), but the lymphatic circuitry involved remained unknown because the anterior and middle fossae of the skull base are difficult to assess by classical immunohistological techniques. In humans, the dural lymphatic architecture remains largely undefined (Absinta et al., 2017; Ding et al., 2021), despite the potential interest of dural lymphatic imaging for the prognosis of neurological diseases.

In this work, we investigated CSF lymphatic drainage with sub-millimeter resolution by large field imaging of the whole head using postmortem light-sheet fluorescent microscopy (LSFM) imaging in mice (Jacob et al., 2019, 2020), and improved real-time magnetic resonance imaging (MRI) in humans (Absinta et al., 2017). Because both techniques preserved the vascular connections between the meninges and the collecting LNs, we were able to establish a 3D-map of the entire lymphatic CSF drainage network in mice and humans. LSFM was performed on decalcified and clarified whole heads after prior administration of a fluorescent tracer into the CSF of mice, while 3T-magnetic resonance (MR) vessel wall imaging (VWI) was carried out in patients after intravenous injection of the low molecular weight contrast agent gadobutrol (Absinta et al., 2017).

Both approaches demonstrated similar circa-cerebral architecture and relationship of MLVs with dural venous sinuses. In both mice and humans, we observed the previously identified dorsal and basal dural lymphatics, and we discovered a large ventral MLV network around the cavernous sinus that includes multiple lymphatic capillary beds and exit circuits through the skull foramina of emissary veins. Remarkably, MRI imaging in humans demonstrated a conserved pattern of cavernous sinus associated MLVs in the anterior part of the brain. We found that cavernous MLVs penetrate the skull through several bilateral foramina of the basal skull (6 and 5 in mice and humans respectively). Notably, we have designed a versatile MRI procedure for mapping of human intra-cranial MLVs that will be highly relevant to the development of diagnostic imaging for patients with CSF drainage defects and neurological diseases.

Collectively, we have demonstrated a spatial organization of the glymphatic outflow and its connection to the dural lymphatic system. The different brain regions, their respective cerebral veins and collecting dural sinuses are in topological relationship with regionally distributed lymphatic capillary beds in the dura mater, thereby providing order to the sampling and drainage of brain antigens toward collecting LNs.

## Results

### LSFM 3D-imaging of cranial CSF outflow pathways

We performed LSFM imaging of whole adult mouse head preparations to visualize cranial CSF outflow (**Fig. 1 A**). Mice were injected with fluorescently-tagged Ovalbumin (OVA-A^555^, 2-8 µl per mouse) into the thoraco-lumbar (Th-Lb) or lumbosacral (Lb-Sa) spinal cord. We chose this site to minimize intracranial CSF pressure increases following injections, and to avoid intracranial tracer leakage that can occur after injections into the cisterna magna. Whole head preparations were decalcified and iDISCO^+^-clarified to allow LSFM imaging through the skull and head tissues (**Fig. 1 A**). OVA-A^555^ spread cranially and caudally from Th-Lb and Lb-Sa injection sites, and its spatio-temporal distribution was similar to other tracers such as india ink or fluorescent FITC-dextran (**Fig. S1, A-G**). OVA-A^555^ deposits were consistently detected along the pia mater and in the perivascular spaces of the spinal cord and brain, as well as within deep cervical LNs (dcLNs) (**Fig. 1, B and C**). A large fraction of the OVA-A^555^ tracer concentrated within phagocytic cells in perivascular spaces of the brain (**Fig. 1 D; Video 1**).

**Figure 1.**
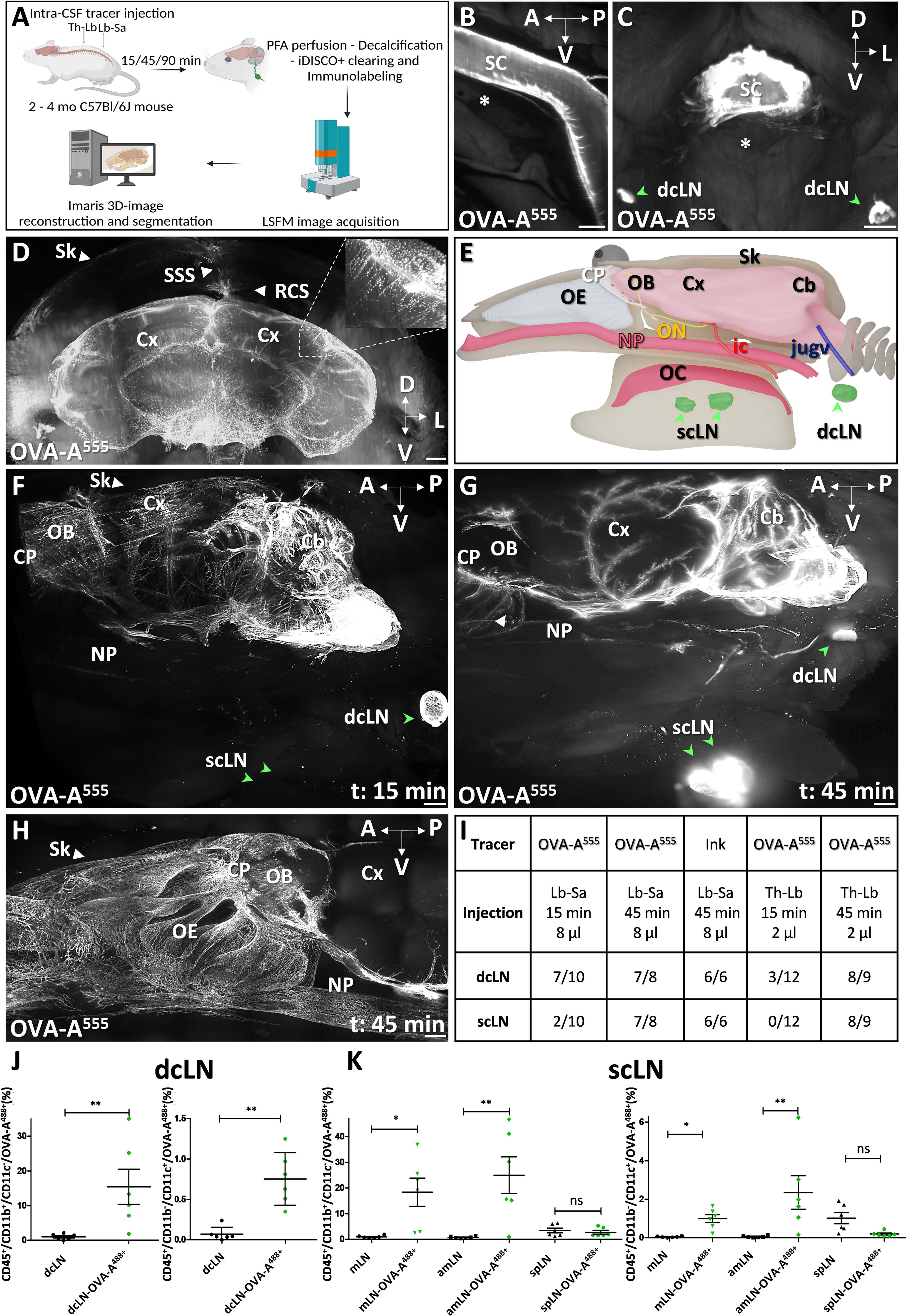
iDISCO-LFSM imaging of cranial CSF tracer drainage. **A** Schematic of the experimental workflow. Gray arrows indicate sites of tracer injection into the spinal cord at the thoraco-lumbar (Th-Lb) or lumbo-sacral (Lb-Sa) level. **B** LSFM imaging of OVA-A^555^ (white) in the pia mater of the spinal cord (SC) 90 min after Th-Lb injection. Asterix: vertebral body. **C** Cross section of cervical SC 45 min after Th-Lb injection shows OVA-A^555^ (white) in the SC and in the deep cervical lymph nodes (dcLN). **D** LSFM view of a coronal section of the forebrain shows OVA-A^555^ distribution around the pia mater, along the perivascular spaces and inside phagocytic cells (inset). Note that brain size is reduced by the iDISCO protocol, but meningeal layers are preserved. The dura and SSS are in contact with the skull (Sk), the pia mater contacts the brain, and tracer drainage can be followed through the intact calvaria. OVA-A^555+^ cells are also found along the superior sagital sinus (SSS) and the rostral confluence of sinuses (RCS). **E** 3D schematic of the lateral view of the head. OB: olfactory bulb, Cx: cortex, Cb: cerebellum, CP: cribrifom plate, ic: internal carotid, jugv: internal jugular vein, OC: oral cavity, OE: olfactory epithelium, ON: optic nerve, dcLN: deep cervical lymph node, NP: nasopharynx, scLN: superficial cervical lymph nodes. **F-H** LSFM lateral views of OVA-A^555^ in decalcificed and cleared whole heads isolated from mice sacrificed 15 min (**F**) or 45 min (**G, H**) after Lb-Sa injection. OVA-A^555^ labeled perivascular g-lymphatic spaces at the brain surface in **F and G**. OVA-A^555^ labeled the dcLN but not the scLN at 15 min (green arrowheads in **F**), while both dcLN and scLN are labeled at 45 min (**G**). Note lymphatic afferent vessels extending from the nasopharynx (NP) towards the LNs (**G**). **H** We observed the same pattern in other mice with a zoom in the anterior part of the CP and the OE. Note a dense plexus of Ova-A^555^+ vessels in the OE (white arrowhead in g), and presence of OVA-A^555^ around the NP, Sk: skull. **I** Summary of tracer injectons and conditions and numbers of tracer-positive cervical LNs per mouse. **J-K** Graphics representing the percentage of OVA-A^555+^ macrophages (CD45^+^/CD11b^+^/CD11c^-^) or dendritic cells (CD45^+^/CD11b^-^/CD11c^+^) among total CD45^+^ cells FACS-sorted from the mandibular lymph node (mLN), the accessory mandibular lymph node (amLN) and the superficial parotid lymph node (spLN) of control mice or mice injected in ThLb with OVA-A^488^. *n*L=L6 mice/group and data show mean+/−SD (error bar) in (**J, K**); Mann–Whitney U test (**J**) and one-way ANOVA with Tukey’s multiple-comparisons test (**K**); *pL<L0.05, **pL<L0.01. ns: non significant. A: anterior, D: dorsal, L: lateral, P: posterior. V: ventral. Scale bar: 500 μm (**B**-**D**,**H**); 800 μm (**F**,**G**).

To study the kinetics of CSF drainage from the meninges to collecting LNs, we examined sagittal views of clarified whole head preparations (**Fig. 1 E-I**). At 15 min after OVA-A^555^ injection, tracer localized in the perivascular spaces of the cortex and olfactory bulbs as well as in dcLNs, but not in the cribriform plate and olfactory epithelium (**Fig. 1 F**). At 45 min after OVA-A^555^ injection, the OVA-A^555^ pattern extended beyond the cribriform plate into the olfactory epithelium, and also labeled the superficial cervical LNs (**Fig. 1 G and H; Fig. S1, H and I; Videos 2 and 3**). CSF drainage kinetics was conserved among mice administered with different volumes of tracers, at different spinal injection sites, and with different tracers (**Fig. 1 I**).

The cellular distribution of tracers in the superficial and deep cervical LNs was analyzed by flow cytometry 90 min after spinal intrathecal injection of OVA-A^488^ (8 μl). Lymph nodes were dissected and labeled with antibodies recognizing macrophages (CD45^+^, CD11b^+^, CD11c^-^) and dendritic cells (CD45^+^, CD11b^-^, CD11c^+^) and sorted by FACS (**Fig. 1 J and K**). OVA-A^488^ expresses a compatible fluorochrome (green) with FACS-sorted labeled LN immune cells. Compared to uninjected mice, the percentage of OVA-A^488+^ macrophages and dendritic cells was significantly increased in dcLNs, as well as in mandibular- and accessory mandibular-scLNs, but not in parotid-scLNs, suggesting that this latter group of cervical LNs is not involved in CSF drainage.

Overall, LSFM imaging after intraspinal tracer injections provided a 3D-view of cranial CSF outflow patterns, showed the rapid uptake of CSF macromolecules by phagocytic cells in perivascular spaces, and revealed multiple circuits of CSF drainage toward cervical LNs.

### Tracer uptake by MLVs in the calvaria and the posterior fossa of the skull base

In all subsequent experiments, OVA-A^555^ (8 µl) was injected into the Lb-Sc spinal cord and mice were sacrificed 45 min later for iDISCO^+^ immunostaining and LSFM imaging of head lymphatic drainage. We established a 3D-map of dural veins and sinuses (insert in the right corner in **Fig. 2 A** using anti-von Willbrand Factor (vWF)-immunolabeling (**Fig 2 B, C, F and I**) overlayed onto the rat dural venous anatomy (Scremin, 2004). MLVs were labeled with anti-LYVE1 (lymphatic vessel endothelial hyaluronan receptor 1) antibody. Tracer deposits and MLVs were then localized with respect to the dural veins.

**Figure 2.**
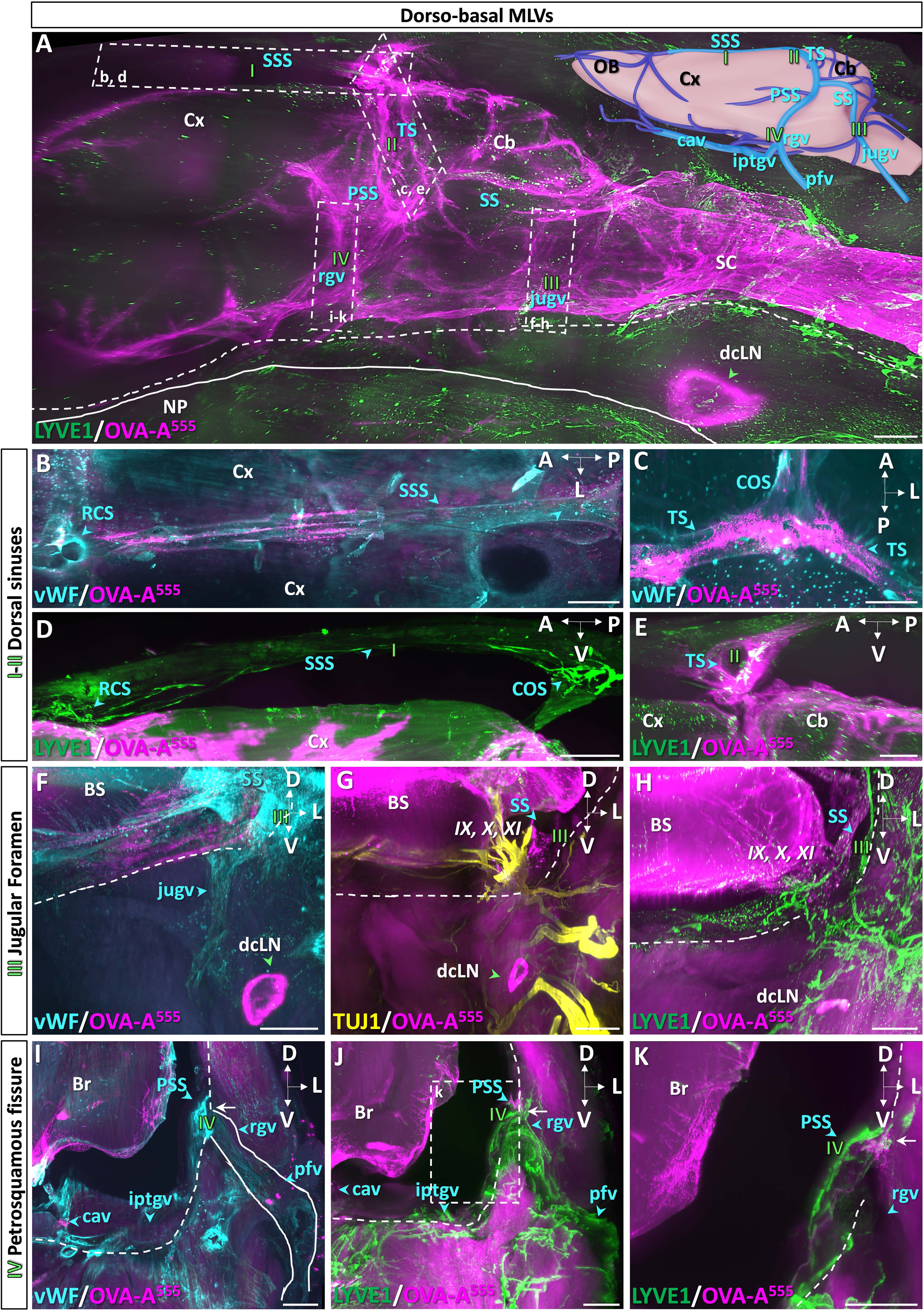
Dorsal and basal meningeal lymphatic drainage. **A** LSFM lateral view of OVA-A^555^ (magenta) in the posterior head after anti-LYVE1 antibody labeling (green). The animal was sacrificed 45 min after Lb-Sa OVA-A^555^ injection. OVA-A^555^ is concentrated in brain meninges, within the perivascular spaces along the spinal cord (SC), the cerebellum (Cb) and the cortex (Cx). Outside of the ventral skull border (dashed line), OVA-A^555^ is detected in the dcLN. NP: nasopharyngeal cavity (solid line). Dashed rectangles correspond to regions magnified in the indicated panels, MLVs and tracer uptake region are numbered in green (I-IV). Inset in **A** shows a schematic of dural veins and sinuses (light and dark blue), the light blue veins are the focus of this figure: cav: cavenous sinus, iptgv: interpterygoid emmissary vein, jugv: jugular vein, pfv: posterior facial vein, PSS: petrosquamous sinus, rgv: retroglenoid vein, SS: sigmoid sinus, SSS: superior sagittal sinus, TS: transverse sinus. **B-E** LSFM horizontal (**B, C**) and sagittal views (**D, E**) of the calvaria, dorsal sinuses are stained with vWF antibody (**B, C**) and MLVs with LYVE antibody (**D**, **E**). The SSS connects the RCS with the COS (**B**, blue arrowheads) which separates in two branches forming the TS (**C**). Note discontinuous OVA-A^555^ labeling in the perisinus spaces (**B**, **C**). LYVE-1+ MLVs line the SSS and TS (**D,** I) and contain OVA-A^555^ at the TS (**E,** II), at the COS and the RCS (**D**), blue arrowheads. **F-H** LSFM coronal views at the level of the foramen of the jugular vein labeled with the indicated antibodies and OVA-A^555^. vWF stained the SS and the jugv, OVA-A^555^ is accumulated around the jugular foramen. (**G**) TUJ1 stained nerves IX, X and XI, which cross the skull through the jugular foramen, OVA-A^555^ is close but not associated directly with the nerves. (**H**) LYVE1^+^ MLVs follow the SS until the jugular vein exits the skull by the jugular foramen. Some MLVs transport OVA-A^555^ positive phagocytic cells (III). *IX*: cranial nerve 9 (Glossopharyngeal), *X*: cranial nerve 10 (Vagus), *X*: cranial nerve 11 (Spinal accessory). **I-K** LSFM coronal views of the petrosquamous sinus exit through the skull.(**i**) vWF stained the PSS passing through the petrosquamous sinus foramen (white arrow) to join the pfv via the rgv, OVA-A^555^ is accumulated around the jugular foramen. (**J**, **K**) LYVE1^+^ MLVs follow the PSS until the rgv and the pfv. Some MLVs show OVA-A^555 +^ phagocytic cells at the petrosquamous fissure level (blue arrowhead white arrow?). Scale bar: 1000 μm (**A**-**D**); 500 μm (**E**-**J**); 250 μm (**K**).

In **Fig. 2 A**, a sagittal view of the caudal part of the head shows the patterns of both OVA-A^555^ deposits and MLVs. Dural sinuses and lymphatics are co-localized in both dorsal (**Fig. 2 B-E**) and basal (**Fig. 2 F-K**) parts of the skull. In the dorsal region, the tracer was discontinuously distributed along the sagittal (**Fig. 2 B**) and transverse (**Fig. 2 C**) sinuses. Tracer accumulated in the peri-sinus spaces as shown around the transverse sinus in **Fig. S2 A and B**. MLVs follow the sagittal and transverse sinuses (**Fig. 2 D and E**). We detected OVA-A^555^ within LYVE1^+^ MLVs in three hotspots at the level of the rostral and caudal confluence of sinuses, and at the transverse sinuses (**Fig. 2 E; Fig. S2 C and D**).

Downstream and toward the basal part of the skull, the transverse sinus extends both caudally into the sigmoid sinus that connects to the jugular vein, and rostrally into the petrosquamosal sinus toward the retrogenoid vein (**Fig. S2 E**). The foramen of the jugular vein, which exits the basal skull along with the *IX*, *X* and *XI* cranial nerves (**Fig. S2 F and G**), displayed a dense network of dural MLVs in contact with the sigmoid sinus, including LYVE1^+^/OVA-A^555+^ capillaries (**Fig. 2 H**). Downstream of sigmoid MLVs, lymphatic vessels exit the skull through the jugular vein foramen and connect to the peripheral lymphatic network that drains into the dcLN (**Fig. 2 F, G and H**). A similar pattern of MLVs was observed on the inner side of the petrosquamous fissure (**Fig. 2 I-K**), in contact with the petrosquasmous sinus (**Fig. 2 K**). Downstream of petrosquamous MLVs, lymphatics exited the skull along the retroglenoid vein, then the posterior facial vein (**Fig. 2 I, J**). In summary, focusing LSFM imaging of lymphatics on previously explored regions of the head confirmed in 3D the MLV pattern and tracer uptake hotspots that had previously been identified on skull cap preparations and sections (Aspelund et al., 2015; Louveau et al., 2015; Antila et al., 2017; Ahn et al., 2019).

### Lymphatic circuits of cavernous sinuses in the middle fossa of the skull base

We then explored lymphatic drainage at the middle fossa of the skull base and the region of the cavernous sinus (**Figs. 3 A**, **4 A and B**). Whole-head preparations were immune-labeled with antibodies recognizing Van Willebrand factor (vWF) to identify veins and with pan-endothelial markers CD31 and podocalyxin (PDLX). We located the cavernous sinus caudally on both sides of the pituitary gland and its connection by the inter-cavernous sinus (**Fig. 3 B; Fig. S3 A-C**). The bilateral cavernous sinuses communicate with the jugular veins via the inferior petrosal sinuses (**Fig. 3C**) and with the transverse sinuses via the superior petrosal sinuses (**Fig. 3 D and E**). They are crossed by the internal carotids (**Fig. 3 A**) and collect blood of the basilar venous plexus and the posterior basal veins (**Fig. 3 C and E**).

**Figure 3.**
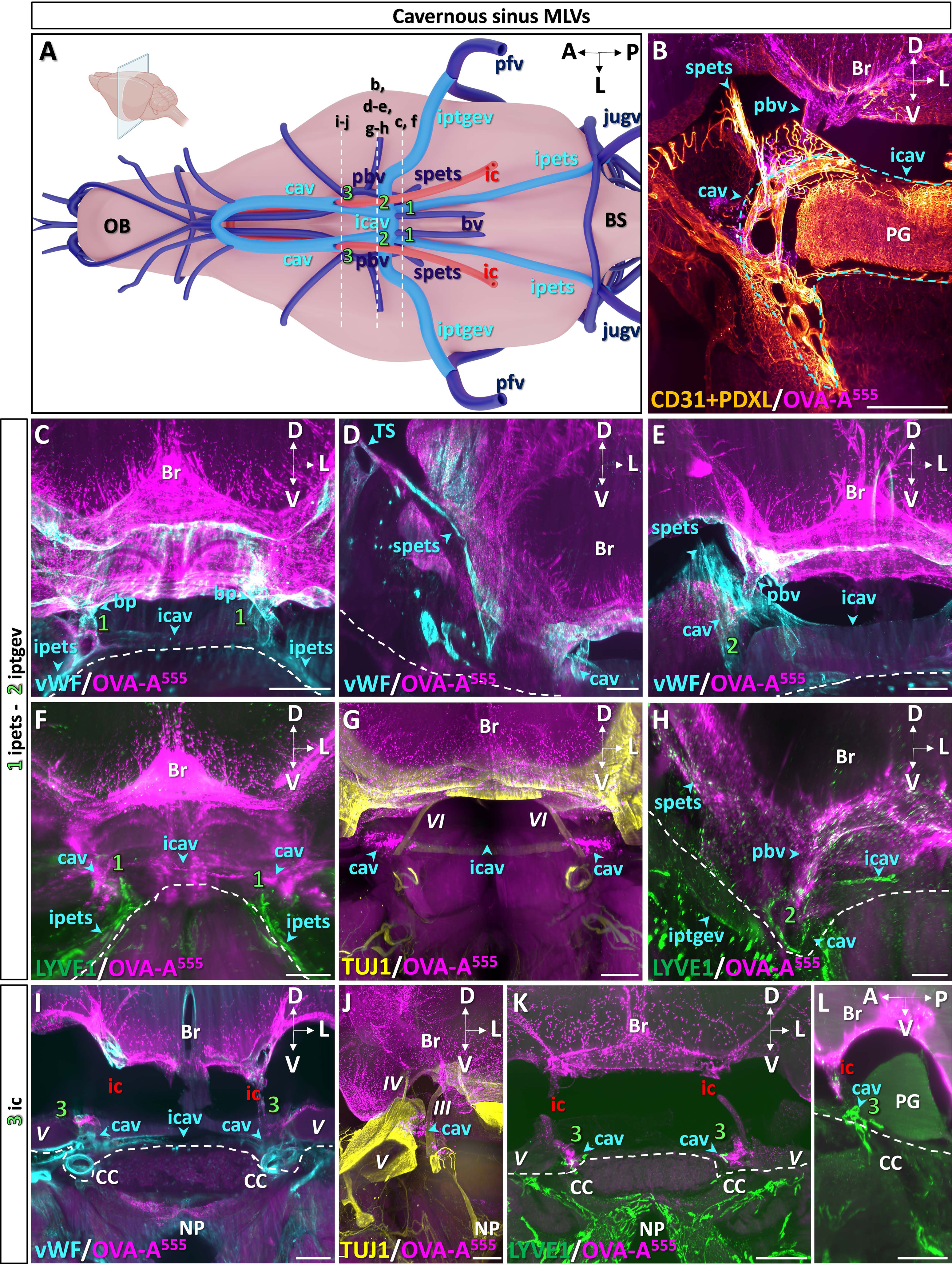
LSFM 3D-images of MLVs associated with the caudal cavernous sinuses in the middle fossa of the skull base. **A** Schematic of the venous sinuses, veins and internal carotid artery at the base of the brain. Stippled areas correspond to the 3 coronal panels (1-3) shown in (**C**, **F**) for 1, (**E**, **H**) for 2 and (**I**, **J**) for 3. Inset in **A**: coronal section plane of the head used in **B**-**K**. bv: basilar venous plexus, cav: cavernous sinus, ic: internal carotid artery, icav: inter-cavernous sinus, ipets: inferior petrosal sinus, iptgev: interpterygoid emmissary vein, jugv: internal jugular vein, OB: olfactory bulb, pbv: posterior basal vein, pfv: posterior facial vein, spets: superior petrosal sinus. **B** OVA-A^555^ was injected into Th-Lb 45 min before sacrifice and sample was labeled with the indicated antibodies. Blood vessels of the pituitary gland (PG) and dura mater express CD31 and PDLX (blue arrowheads). Tracer deposits (magenta) were observed in the pia mater of the brain, around the cavernous sinus (cav) and the posterior basal vein (pbv). **C-E** Ventral (**C**, **D**) and dorsal (**E**) views of a sample stained with vWF to label dural sinuses and veins (blue arrowheads). TS: transverse sinus. Tracer accumulated in peri-venous and -sinusal spaces, especially at confluence points. Dashed line: limit between intracranial and extracranial regions. **F-H** Lymphatic vasculature (LYVE1^+^, green in **F**, **G**) and TUJ1^+^ cranial nerves (yellow in **G**), *VI*: cranial nerve 6 (Abducens) at coronal levels 1 and 2. MLVs contacted the cavernous perisinus space where tracer accumulated (**F**), and surrounded the foramen of iptgev (**H**). Cranial nerves were devoid of tracer deposits and MLVs. **I-L** Dural veins (vWF, blue in **I**), lymphatic vasculature (LYVE1^+^, green in **K**, **L**) and cranial nerves (TUJ1^+^, yellow in **J**) on coronal (**I-K**) and sagittal (**L**) views at level 3. MLVs contacted the cavernous sinuses (**K**, **L**) and lymphatic tracer uptake (white in **K**) was detected at the intersection of cavernous sinuses with internal carotid arteries in the skull base. Cranial nerves showed neither tracer deposits nor MLVs (**J**). *V*: cranial nerve 5 (Trigeminal), *IV*: cranial nerve 4 (Trochlear), *III*: cranial nerve 3 (Oculomotor). Br: brain, CC: carotid canal, NP: nasopharyngeal cavity. A: anterior, D: dorsal, L: lateral, P: posterior, V: ventral. Scale bar: 500 μm (**B**-**L**).

**Figure 4.**
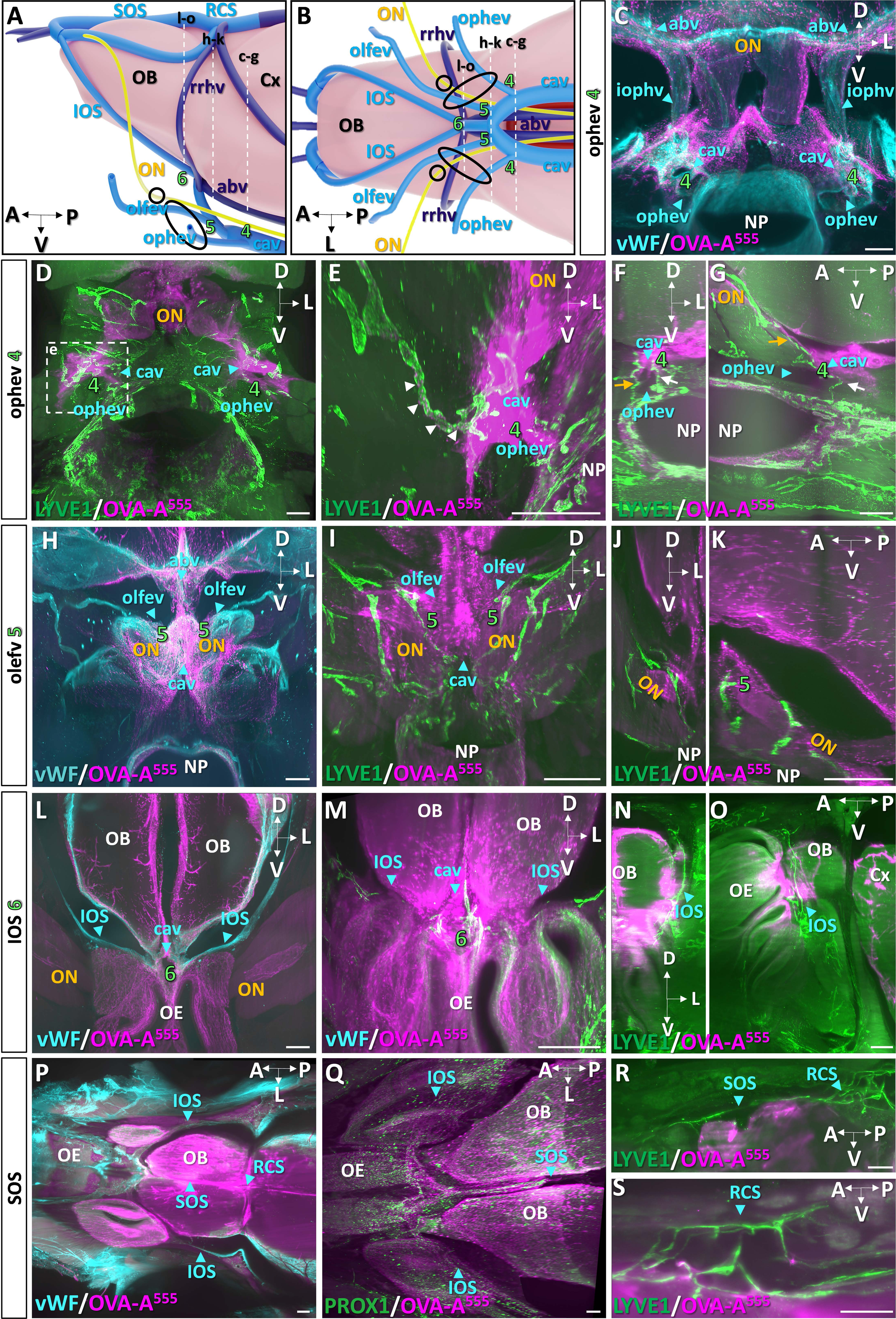
Rostral cavernous sinus drainage by MLVs and CSF outflow into the orbital cavity and NP. **A, B** Schematics of the meningeal vasculature of the forebrain and olfactory bulbs. Lateral (**A**) and ventral (**B**) views. Stippled areas correspond to the 3 coronal panels (4-6) shown in (**C-G**) for 4, (**H-K**) for 5 and (**L-O**) for 6. Black circles: optic nerve foramen and anterior lacerated foramen (of emissary veins). abv: anterior basal vein, Cx: cortex, olfev: olfactory emissary vein, ON: optic nerve (yellow), ophev: ophthalmic emissary vein, rrhv: rostro-rhinal vein. **C** Coronal view of cerebral veins/sinuses (vWF^+^, blue) and tracer deposits (OVA-A^555^, magenta) in perivenous (internal ophthalmic vein (iophv)) and perisinus (cavernous sinus) spaces at the level of the ophthalmic emissary vein exit (ophev) (4). **D-G** MLVs (LYVE1^+^, green) and tracer deposits (magenta) located around the cavernous sinuses (green, 4). Coronal (**D-F**) and sagittal (**G**) views. MLV capillary directly contacting peri-sinus deposits (**D, E**) and transporting OVA-A^555+^ phagocytic cells (white arrows in **e**, magnification of **D**). OVA-A^555+^ MLVs also follow the ophthalmic emissary vein to exit the skull through the anterior lacerated foramen toward the orbital cavity (orange arrows in **G**). Lymphatic drainage of tracer was also detected along the nasopharynx (NP, white arrows). **H-K** Menigeal veins (vWF, blue in **H**), lymphatic vasculature (LYVE1^+^, green in **I-K**) and tracer deposits (OVA-A^555^, magenta) on coronal (**H-J**) and sagittal (**K**) views at level 5. Tracer accumulated around the anterior basal and rostro-rhinal veins and the cavernous sinuses (**H**). Tracer deposits were found in the pia mater of the optic nerve (**I-K**), but lymphatic tracer uptake was only detected around the cavernous sinus (white in **I**) and not along the optic nerve inside the orbital cavity (**I**). **L-O** Rostral end of cavernous sinuses at level 6. Dural veins (vWF, blue in **L**), lymphatic vasculature (LYVE1^+^, green in **M-O**) and tracer deposits (OVA-A^555^, magenta). MLVs of the inferior olfactory sinus (IOS) (**M**, **N**, **O**, blue arrowhead) and lymphatic tracer uptake at the rostral end of the cavernous sinus (white in **M**). Coronal (**N**) and sagittal (**O**) views of MLVs running from the peri-sinus cavernous area to the olfactory bulb (**N, O**). **P-S** Horizontal views of the anterior part of the head showing the inferior and superior olfactory sinuses (IOS and SOS, respectively) and the rostral convergence of sinuses (RCS). Dural veins (vWF, blue in **P**), lymphatic vasculature (green in **Q-S**) and tracer deposits (OVA-A^555^, magenta). MLVs are labeled with either anti-PROX1 (**Q**) or anti-LYVE1 (**R, S**) antibodies. MLVs connect the cavernous sinuses with the rostral convergence of sinuses via olfactory sinuses. Scale bar: 300 μm (**C**-**S**).

We identified multiple lymphatic foci associated with the cavernous sinuses, as schematized in **Fig. 3 A** and **Fig. 4 A and B**. In the caudal region of cavernous sinuses, MLVs localized parallel to the inferior petrosal sinuses (**Fig. 3 F**), as well as at the foramen of the inter-pterygoid emissary veins and along the inter-cavernous sinus (**Fig. 3 H**). LYVE1^+^/OVA-A^555+^ capillaries were concentrated at the intersection of the cavernous sinuses with internal carotid arteries (**Fig. 3 I-L**; **Video 4**). In contrast, OVA-A^555^ deposits were not detected along neighboring cranial nerves (*VI* in **Fig. 3 G** and *III*, *IV* and *V* in **Fig. 3 J**). At the confluence of ophthalmic and olfactory emissary veins with the rostral part of cavernous sinuses (**Fig. 4 A-C and H; Fig. S4 A-C and E**), the OVA-A^555+^ peri-sinus area harbored bilateral LYVE1^+^/OVA-A^555+^ foci (**Fig. 4 D, E and L; Fig. S4 D and F; Video 5**). These lymphatic capillaries showed circulating OVA-A^555^-loaded phagocytic cells which attests the cell-mediated transport of CSF macromolecules in MLVs (white arrowheads in **Fig. 4 E**). LYVE1^+^/OVA-A^555+^ vessels of the cavernous sinus exited the skull by the anterior lacerated fissure toward the lymphatic beds of the orbital cavity and the nasopharynx (**Fig. 4 F, G, J and K**). OVA-A^555^ also labeled the pia mater and arachnoid layers all along the optic nerve until the retina (**Fig. 4 C; Fig. S4 G and H).** In the most rostral part of the cavernous sinus, other LYVE1^+^/OVA-A^555+^ capillaries contacted the inferior olfactory sinuses and were prolonged by dural lymphatics running along the superior olfactory sinus toward the dorsal surface of olfactory bulbs (**Fig. 4 L-O; Fig. S4 I-K**), and the rostral confluence of sinuses (**Fig. 4 P-S**).

### Facial lymphatic drainage from ethmoidal and orbito-nasal regions

The lamina cribrosa of the ethmoid bone, also called the cribriform plate, is a main exit pathways of CSF outflow from the mouse skull (Norwood et al., 2019). It remains unclear if MLVs contribute to this process and extend through the cribriform plate toward the nasal cavity (Proulx, 2021). As shown on a lateral view of the forebrain and the nasal cavity (**Fig. 5 A**), we found OVA-A^555^ along perivascular spaces of the cortex and olfactory bulbs, exiting through the cribriform plate, and downward into the olfactory and respiratory epithelia. In the ethmoid bone dura, LYVE1^+^ capillary-like lymphatics were observed close to the olfactory perivascular spaces and the cribriform plate (**Fig. 5 B**). These ethmoid MLVs concentrated between the olfactory bulbs, surrounding the olfactory nerve foramina, and they colocalized with a dense population of LYVE1^+^/OVA-A^555+^ phagocytic cells (**Fig. 5 B and C**). Unlike other MLVs, ethmoid MLVs were not associated with dural vWF^+^ sinuses (**Fig. 5 D**). We failed to detect extension of ethmoid MLVs across the ventral and central part of the cribriform plate toward the nasal cavity (**Fig. 5 B; Video 6**). In contrast, we found a rostral lymphatic connection between ethmoid MLVs and the nasal cavity via the foramen cecum, in the most dorso-rostral part of the cribriform plate (**Fig. 5 and E**). MLVs have therefore only a minor contribution to nasal CSF outflow at the cribriform plate.

**Figure 5.**
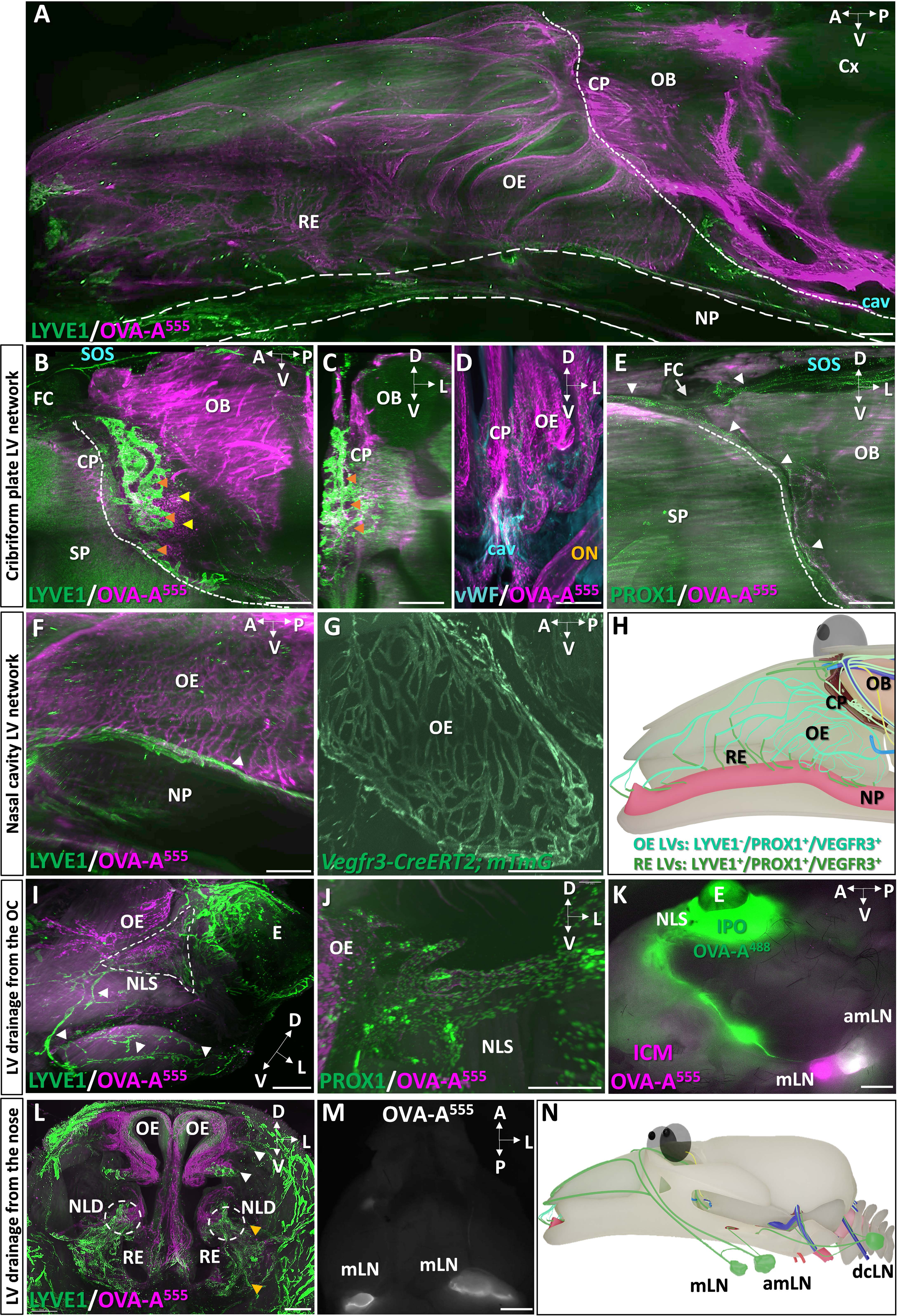
Rostral drainage of CSF from cribriform plate towards to nasal cavity. **A** Sagittal view of the nasal cavity and forebrain of a mouse sacrificed at 45 min after spinal intrathecal injection of OVA-A^555^ tracer. Lymphatics (LYVE-1, green), tracer (magenta). Tracer deposits were detected in meningeal perivascular spaces, the cribriform plate and throughout the olfactory (OE) and respiratory (RE) epithelia of the nasal cavity. Dotted line: limit between intra and extracranial regions; dashed lines: nasopharynx (NP). **B-E** Cribriform plate (CP) region. Lymphatic vasculature (green in **B, C, E**), meningeal veins (vWF, blue in **D**) and tracer deposits (magenta) on sagittal (**B**) and coronal (**C-E**) views of the CP. MLVs were located in the dura of the ethmoid bone and around CP foramina (orange arrowheads in **B**, **C**). No LYVE1^+^ vessels were found to cross the CP toward to nasal epithelia (**B**, **E**), while the outflow of tracer filled the OE and RE (**A, D**). Tracer-labeled phagocytic cells concentrated in the CP dura mater (yellow arrows in **B**). Lymphatic capillaries (LYVE-1^+^) surrounding CP foramina were not associated with a venous sinus (orange arrowheads in **D**). PROX1^+^ lymphatics of the nasal cavity (white arrowhead in **E**) connected with both ethmoid MLVs via the foramen cecum (white arrow) and MLVs of the superior olfactory sinus (SOS). **F, G** Nasal cavity lymphatic drainage. Sagittal views of the nasal cavity showing: the tracer-labeled LYVE-1^-^ vasculature of the nasal epithelium (magenta in **F**), the LYVE1^+^ lymphatics of basal respiratory epithelium and nasopharynx (green in **F**), and the *Vegfr3*-expressing vessels of the nasal epithelia (GFP reporter, green in **G**). **H** Sagittal schematic of olfactory and nasal lymphatic circuits. Olfactory bulb MLVs (fluorescent green) do not cross the CP with olfactory nerves but exit rostrally into the nasal cavity, via the foramen cecum. Olfactory epithelium lymphatics (blue, PROX1^+^/VEGFR3^+^/LYVE1^-^) transport CSF likely collected from perineural drainage along olfactory nerves. They collect into LYVE1^+^ vessels (green) of the basal respiratory epithelium and nasopharynx which connect with cervical LNs. **I-K** Lymphatic drainage from the orbital cavity. Tracer deposits (OVA-A^555^, magenta) and lymphatic vasculature (green in **I, J**). (**I**) Ventrolateral view of the nasal lacrimal sac region (NLS, dashed line) shows tracer deposits between the orbital cavity and the olfactory epithelium (OE). Orbital LYVE1^+^ lymphatics connect with facial lymphatics (white arrowheads). (**J**) Lymphatic tracer uptake by orbital PROX1^+^ vessels (white arrowhead). (**K**) Ventrolateral view of the forehead from a mouse administered with spinal intrathecal OVA-A^555^ (magenta) and a subsequent intra-orbital delivery of OVA-A^488^ (green). Both OVA-A^488^ and OVA-A^555^ collected into the associated-mandibular LNs (white), but OVA-A^555^ only drained into the mandibular LNs (magenta). **L-N** Lymphatic drainage from the nose. (**L**) Coronal view of the nose showing two drainage circuits: dorsal (white arrowhead) and ventral (yellow arrowhead). Dashed circles show the distal part of NLD with surrounding LVs co-stained with OVA-A^555^ (magenta) and with antibody against LYVE1^+^ (green). (**M**) Macroscope image of the ventral forehead showing nasal lymphatics transporting tracer toward mandibular LNs (amLN, mLN). (**N**) Sagittal schematic of lymphatic drainage circuits from the orbital cavity and the nose toward cervical LNs. A: anterior, amLN: accessory mandibular LN, cav: cavernous sinus, CP: cribrifom plate, Cx: cortex, D: dorsal, dcLN: deep cervical L, FC: foramen cecum, L: lateral, mLN: Mandibular LN, NLS: nasolacrimal duct, NP: nasopharynx, OB: olfactory bulb, OE: olfactory epithelium, P: posterior, SP septum, V: ventral. Scale bar: 500 μm (**A**-**J**, **L**); 80 μm (**D**); 2 mm (**K**,**M**).

In the superior and middle part of the nasal cavity, downstream of the cribriform plate, the absence of LYVE-1 lymphatics (**Fig. 5 F**) contrasted with the presence of a dense network of VEGFR3^+^ (**Fig. 5 G; Fig. S5 A**) and Prox1^+^ (**Fig. S5 B**) expressing capillaries that drained OVA-A^555^ phagocytic cells. In the basal domain of the olfactory epithelium and along the nasopharynx, LYVE1 expression reappeared in OVA-A^555^-labeled vessels (**Fig. 5 F**), which are likely collector lymphatics located downstream of the VEGFR3^+^/PROX1^+^/LYVE1^-^ narrow vessels of the upper nasal region. A continuous LYVE1^+^ lymphatic network was observed between these two types of vessels at the contact of the basal olfactory epithelium (**Fig. S5 C**). Downstream of the nasal circuit, the nasopharynx provided a rostrocaudal route for CSF/ISF-draining lymphatics toward cervical LNs (**Fig. 5 F**). Altogether, these observations indicate that lymphatics of the upper nasal cavity are distinct from MLVs, without physical continuity and phenotype similarity, although both lymphatic circuits contribute to CSF drainage (**Fig 5 H**).

Finally, we focused on the more distal pathways of OVA-A^555^ outflow and lymphatic drainage in the orbital and nasal cavities. First, in the ocular region, we found OVA-A^555^ deposits in the nasolacrimal sac area in PROX1-expressing LVs (**Fig. 5 I and J**). Nasolacrimal LYVE1+ LVs collected directly into the associated-mandibulary LNs. As shown in **Fig. 5 K**, after intraorbital delivery of OVA-A^488^ (green) into mice with prior intraspinal OVA-A^555^ injection (magenta), OVA-A^488^ collected exclusively in the associated-mandibulary LNs (white), and not in mandibular LNs (magenta). Second, in the nose, we identified two foci and drainage circuits localized in the dorsal and ventral aspects of the nasal cavity (arrows in **Fig. 5 A**). LYVE1 LVs co-localized with OVA-A^555^ deposits and OVA-A^555^ LYVE1^+^ lymphatics were detected in the basal respiratory epithelium and around the distal part of the nasolacrimal duct (yellow arrowheads in **Fig. 5 L**, magnification in **Fig. S5 D**). We showed that mandibular LNs directly collected fluids from the anterior nasal cavity as they were labeled 5 min after OVA^488^ inhalation (**Fig. 5 M**).

Therefore, both intraorbital and nose LVs collect the CSF/ISF outflow from the cribriform plate. These orbital and nasal lymphatic circuits collect into the associated-mandibular and mandibular LNs, respectively (**Fig. 5 N**).

### Caudal lymphatic drainage of CSF toward vertebral LNs

We examined the caudal outflow of CSF in the sacral region of the spinal cord. Here, we performed i.c.m OVA-A^555^ injection and mice were sacrificed 90 min later as previously reported (Ma et al., 2019). Under a binocular microscope, OVA-A^555^ deposits were observed in the intravertebral spaces of the coccygeal and sacral vertebral column, between S3 and Co2, as well as in the sciatic, lumbar and renal LNs (**Fig. S6 A and B**). LSFM imaging confirmed that OVA-A^555^ accumulated in the dural and subarachnoid space all along the spinal cord (arrow), and specifially in the epidural space between S3 and Co2 (**C-E**). Paraffin sections of sacral vertebral segments isolated after i.c.m. injection of Indian ink revealed a thinning of the dorsal dura mater and clusters of ink-labeled phogocytic cells, likely macrophages, in the epidural S3-Co2 region (**Fig. S6 G and K**).

### 3D-mapping of the dural blood-lymphatic vasculatures in humans

We investigated the vasculature of human meninges to determine if humans and mice displayed a convergent organization of dural lymphatic vasculature. To this aim, MRI was conducted on patients with either dural sinus stenosis (*n* = 5) or multiple sclerosis (*n* = 1). For vascular imaging, the patients received a systemic injection of gadobutrol, just before MRI and the scans were next analyzed using a 3D-image reconstruction software (**Fig. 6 A**). The flow of the contrast agent was imaged sequentially, first in the blood vessels, then after it reached the lymphatic vasculature. In brief, we first used a MR elliptic venography sequence then, at 8 to 12 min after gadobutrol injection, a 3D T1 SPACE (variable flip angle turbo spin echo) VWI sequence modified by addition of a DANTE (delay alternating with nutation for tailored excitation) module (Siemens, Healthineers). The T1 SPACE DANTE sequence allowed to accurately segregate the slow-flow circuits of the lymphatic vessels from the faster flow circuits of arteries, veins, venules, and CSF in a 6-minute scanning time. The combination of the elliptic venography and T1 SPACE DANTE sequences resulted in large field and sub-millimeter resolution images of the blood and lymphatic vasculatures in the meninges and the neck regions (**Fig. 6 B and C**). A 3D-map of the different flow circuits of gadobutrol was finally generated from native sequences, including venous sinuses and cerebral veins, cerebral arteries, and slower flow circuits. As shown in **Fig. 6 D**, the slower-flow circuit (yellow) was concentrated in the peri-sinus areas, standing apart from venous sinuses and veins (blue), and included vessel-like compartments (arrows) associated with flattened vesicles (arrowheads).

**Figure 6.**
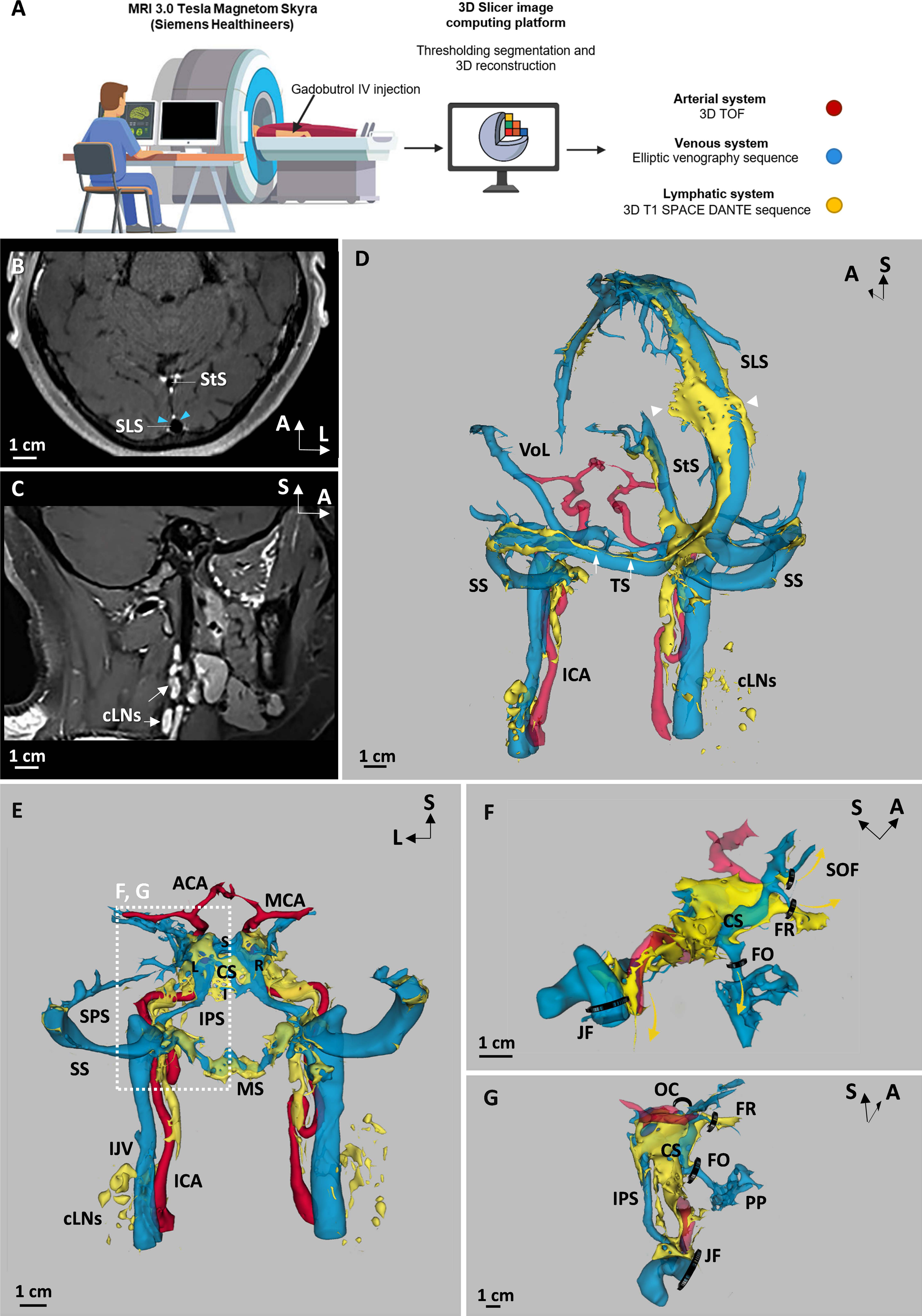
Acquisition and post-processing of meningeal vascular MR-Imaging in humans. **A** Schema of the workflow for meningeal vascular 3D-mapping by MRI in humans. Native sequences were acquired from a 3T Skyra MRI (Siemens Healthineers) before and after intravenous (IV) Gadobutrol injection. 3D-Slicer platform was next used for semi-automatized signal intensity-based thresholding and segmentation of native sequences. 3D TOF (Time of Flight), Elliptic Venography after gadobutrol injection and 3D T1 SPACE DANTE after gadobutrol injection sequences were used to segment the arterial, venous and lymphatic system, respectively. **B** Native 3D T1 SPACE DANTE sequence after Gadobutrol injection in an oblique axial plane crossing the transversal axis of both the superior longitudinal (SLS) and the straight sinuses (StS). The DANTE-improved black blood effect allowed to darken the lumen of venous sinuses, while dural lymphatics (white) were strongly enhanced by the contrast agent and precisely segregated from the unenhanced dura-mater of the parasinus (light blue arrowheads). **C** Native 3D T1 SPACE DANTE sequence after Gadobutrol injection in a sagittal plane along the longitudinal axis of the left IJV covering both the head and neck. The dcLNs (arrows) surrounding the IJV were enhanced by the contrast agent and distinct from adjacent soft tissues. **D.** Oblique-posterior view of the dorsolateral group of dural venous sinuses (blue), including the SLS, the StS, the transverse (TS) and the sigmoid sinuses (SS). The vein of Labbé (VoL) represents the major venous inflow of lateral sinuses (TS + SS) at a confluence point located between the transverse and the sigmoid sinuses. The IJV represents the major venous outflow of the dorsolateral group of sinuses. The related dorsolateral parasinus fluids (yellow) are concentrated in peri-sinus areas and include vessel-like compartments (arrows) associated with flattened vesicles (arrowheads). A significant perivascular route of parasinus fluid was detected along the internal carotid arteries (ICA) and join the dcLNs. **E** Posterior view of the meningeal vascularization in the anterior part of the skull. Left (L) and right (R) cavernous sinuses are connected at the midline by the superior (S) and inferior coronary sinuses (I). The superior petrosal sinus (SPS) runs in the attached margin of the tentorium cerebelli and connects the CS with the SS, while the inferior petrosal sinus (IPS) connects the CS with the IJV. The IPS is also connected with the marginal sinus (MS) which drains caudally in the peri vertebral venous plexuses. In their intra-cavernous segments, the internal carotid arteries (ICA) cross the CS before their intradural bifurcation in middle (MCA) and anterior (ACA) cerebral arteries. Parasinus fluids were detected in the para-sinus areas of the CS, the MS surrounding the foramen magnum, as well as along the IPS. The exit routes of parasinus fluids from the skull followed the peri-carotid route in the carotid canal, anteriorly, and the peri-vertebral canal along the vertebral arteries, posteriorly. **F, G** Exit routes of CS parasinus fluids. The parasinus flow around the CS (arrows) followed the peri-carotid route inside the carotid canal as well as several trans-foraminal routes along the trigeminal nerve (V) branches including the superior orbital fissure (SOF) along the ophthalmic branch (V1), the foramen rotundum (FR) along the maxillary branch (V2), and the foramen ovale (FO) along the mandibular branch (V3). Besides their corresponding trigeminal nerve branches, these foramina also contained their corresponding emissary veins which drained extracranially in the pterygoid plexuses (PP). No parasinus flow was observed through the optical canal (OC). Scale bar: 1cm (**B**-**G**).

### Circa-dural lymphatic drainage in humans

We have tracked the slower flow of gadobutrol throughout the human dura. VWI scans were focused along all dural venous sinuses in the dorsolateral, anterior and posterior regions of the skull, where we had previously mapped dural lymphatics in the mouse. The presence of dorsolateral circuits of drainage was confirmed along the superior sagittal, straight, transverse, and sigmoid dural sinuses (**Fig. 6 D**), in agreement with previous reports (Absinta et al., 2017). Interestingly, parallel to the superior sagittal sinus, we found perforating venules and slow flow channels crossing the skull and connecting to superficial intra-calvaria and subcutaneous lakes (**Fig. S7 A**). Connection of dural channels with calvaria bones has been previously observed in the mouse (Cai et al., 2019; Pulous et al., 2021), and only recently in humans (Ringstad and Eide, 2021). In the anterior region of the skull, the slow flow of gadobutrol was detected in the para-sinus area of the cavernous sinus (**Fig. 6 E-G**). In the inferior region of the skull, the marginal sinus around the foramen magnum was also surrounded by gadobutrol slow flow which extended caudally along the vertebral arteries (**Fig. 6 E**). Like dural lymphatics in the mouse, the regional circuits of gadobutrol slow flow were interconnected. Slow flow of gadobutrol was observed along the inferior petrosal sinus, which links the cavernous sinus with the jugular bulb, therefore connecting the anterior and dorsolateral regions (**Fig. 6 E-G**). Moreover, the anterior, and dorsolateral regions of the skull were connected with the inferior region along the marginal sinus that connects the jugular bulb and the internal petrosal sinus with the peri-vertebral venous plexuses (**Fig. 6 E**).

The exit of lymphatics from the skull was observed along blood vessels through the jugular foramen for dorsolateral lymphatics, and along the carotid canal, the superior orbital fissure, the foramen rotundum, and the foramen ovale for lymphatics of the cavernous sinus (**Fig. 6 F and G**). In contrast to caudal regions, the most frontal region of the skull was devoid of lymphatic vessels and, as in mice, we failed to detect lymphatic connection with the nasal conchae through the central part of the cribriform plate (**Fig. S7 B**). In the neck region, extracranial lymphatics were observed in continuity with dural lymphatic vessels and also in connection with cervical lymph nodes (**Fig. 6 D**).

Therefore, the circuits of peri-sinus drainage mapped by MRI in the human dura mater (**Fig. S7 C**) mirror the dural lymphatic circuits identified by LSFM in the mouse (**Fig. S5 G**). This convergence of data leads us to propose that the distribution pattern of lymphatic vessels in the human dura can be predicted from the gadobutrol slow flow detected by T1 SPACE DANTE sequences.

## Discussion

We have tracked lymphatic drainage in the dura mater using two procedures of tracer delivery, either the intra-spinal administration of ovalbumin in mice, or the systemic injection of gadobutrol in humans. Both approaches resulted in a similar pattern of tracer around dural venous sinuses and in cervical LNs. In both species, we found that the venous sinuses were associated with lymphatic vessels, including the tracer-containing capillaries, in all regions of the dura mater. These findings indicate a specific role of dural lymphatics in the uptake and drainage of the glymphatic efflux from para-sinus areas. This dural lymphatic uptake may conveniently preserve the subarachnoid CSF from pollution by the brain-derived waste transported in the perivascular space of cerebral veins.

In mice, fluorescent macromolecule tracer was detected in phagocytic cells located along brain perivascular spaces, in the peri-sinus area and within dural lymphatics at hotspots where cerebral veins enter venous sinuses. This observation supports a model of phagocytic uptake and phagocytic cell transport of macromolecules from the perivascular spaces of cerebral and dural veins into the dural lymphatics. Dural lymphatic function would thus be, at least partially, dedicated to the sampling and transport of glymphatic fluids and to the phagocyte-mediated transport of peri-sinus macromolecules. Additional drainage functions of dural lymphatics at peri-sinus hotspots may also include the direct uptake of CSF from the subjacent subarachnoid space or the surrounding dural tissues. The perivascular lymphatic pathway is considered as a main route for the drainage of antigens from the brain to cervical LNs (Engelhardt and Coisne, 2011) but it remains unclear whether and how perivascular myeloid cells can traffic via this route to the dura mater and thereby mediate neuroimmune communication. Under physiological conditions, meningeal and perivascular macrophages are known to be dynamic cells that sample their environment (Nayak et al., 2012, 2014). Macrophages and dendritic cells are also present in the dura mater, but the origin and dynamics of these cells remain discussed (Goldmann et al., 2016; Prinz et al., 2017; Van Hove et al., 2019; Rua and McGavern, 2018). Our present localization of glymphatic efflux hotspots in the dura mater may facilitate the dynamic imaging of perivascular and dura mater myeloid cells and help to better understand their role in brain antigen export into the peripheral immune system.

LSFM imaging of the mouse head allowed us to map tracer deposits together with lymphatic circuits, from uptake hotspots in the dura to collector vessels and cervical LNs. This postmortem approach to map CSF drainage, however, required extensive incubations into buffers that wash tissues and eliminate part of the fluorescent tracer injected into the CSF. Thus, the LSFM map of CSF drainage may not be exhaustive and may not account for the rapid bulk flow of CSF that is known to privilege perineural routes toward collecting LNs (Engelhardt and Coisne, 2011). The VWI technique provided us with an alternative realtime visualization of dural fluid drainage, by setting apart lymphatics from blood vessels according to their different flow rate. Although this approach lacks a histological assessment of flow circuits, its accuracy to map dural lymphatic drainage in humans is supported by the convergence of VWI and LSFM imaging data, independent of species identity.

The 3D-maps of dural lymphatics in mice and humans highly overlap throughout the dorsal, lateral, anterior and posterior regions of the skull. In addition to the dorsal and lateral lymphatics already identified in the calvaria and the posterior fossa of the skull base, we discovered a large lymphatic bed around the cavernous sinus that connected with dorsal and basal lymphatics. Lymphatic drainage from the cavernous sinus area was mapped along the trigeminal nerve branches and through the foramina and fissures of the anterior skull. Cavernous lymphatics may thus drain peri-venous efflux from their tributary cerebral veins (superficial and deep middle cerebral veins, and hypophyseal vein), and possibly from orbital veins (superior and inferior ophthalmic), into collecting scLNs and dcLNs. The nasopharynx provided a support to the most anterior lymphatic drainage from the cavernous sinus, in agreement with the recent observation that the nasopharyngeal route has a main role in the extracranial drainage of CSF outflow (Decker et al., 2021).

In the mouse, we failed to find a connection between ethmoid MLVs and the lymphatic vessels of the nasal cavity. A striking contrast was observed between the phenotypes of olfactory MLVs (LYVE-1^+^ capillaries) and lymphatics of the upper/middle nasal epithelium (narrow LYVE-1^-^ vessels). Although both MLVs and nasal lymphatic circuits drained CSF tracers, we propose that they form distinct lymphatic beds endowed with different drainage function, mainly dedicated to the glymphatic outflow for MLVs and to the uptake of subarachnoid CSF for nasal lymphatics. In humans, the contribution of lymphatics to cribriform plate drainage was also not supported by VWI data. Gadobutrol flow was not detected through the cribriform plate, but it did accumulate in the nasal cavity. Nevertheless, we cannot rule out the possibility that thin lymphatics, below the resolution limit of VWI detection, can contribute to cribriform plate drainage in humans. Although highly overlapping in their spatial distribution, dural lymphatics in mice and humans showed different morphologies. MLVs are thin in mice while they are associated with large sacs of peri-sinus fluid drainage in humans. MRI data have been only collected so far from patients with possible alterations of MLVs, including bilateral assymetries or irregularities. We will thus need further MRI and additional LSFM imaging of the human dural vasculature, especially in controls without vascular stenosis or neurological diseases, to definitively establish the structural and functional relationships between lymphatic vessels and other peri-sinus non-vascular structures involved in dural fluid drainage.

Dural lymphatic vessels were imaged in humans, for the first time, using VWI techniques and 3D T1 SPACE sequence (Absinta et al., 2017). We combined this imaging technique with MR venography to unanambiguously discriminate lymphatics from venous sinuses and veins. The 3D T1 SPACE sequence is also limited by an incomplete suppression of slow flow that may lead to small venous vessels being misidentified as dural lymphatics (Bapst et al., 2020). Accordingly, we have incorporated the DANTE module in the 3D T1 SPACE sequence to improve the suppression of the residual slow-flow signal and optimize the black-blood effect. Therefore, this MR-imaging protocole allows a routine exploration of the whole brain and neck vasculature, including lymphatics, in a timely, reliable and accurate manner, which is especially attractive in clinical practice.

The 3-D map of the veno-lymphatic complex highlights the close anatomical relationship existing between the lymphatics and the peri-sinus and -venous areas in the dura mater (**Fig 6 D and E**), a common feature in humans and mice. This anatomical evidence suggests that the neuro-immune interface initially described in the peri-sinus area along the superior sagittal sinus (Rustenhoven et al., 2021) likely exists around most venous sinuses of the skull, at least in the mouse. The different lymphatic hotspots around the dura mater sinuses may allow the sampling of region specific-information from the brain, thereby confering a regional specificity to the immune surveillance of the CNS in the meninges.

In humans, the pattern of cortical veins can vary between individuals and, for each person, between hemispheres (Kılıç and Akakın, 2008). As a consequence, the patterns of peri-venous glymphatic efflux and dural lymphatic uptake may vary between individuals and pathophysiological conditions. Remodeling processes are thus predictable at dural lymphatic uptake sites and thoughout CNS lymphatic drainage circuits in humans, especially when considering the rich inter-connections between dural sinuses and the well-known phenotypic plasticity of the venous and lymphatic systems. The functional predominance of one regional dural lymphatic system over another one, for example, the dorso-lateral over the anterior system, may change upon cooperative or competitive regulatory mechanisms that remain to be determined. Otherwise, exploration of the variability and plasticity of the veno-lymphatic complex should provide new insights into the region-specific vulnerability of neurodegeneration or inflammation observed in humans.

The accuracy of T1 SPACE DANTE sequence moreover allowed us to detect veinules and small non-blood vessels that penetrate together through the skull and join calvarian lakes located parallel to the superior sagittal sinus (**Fig. S7 C**). These trans-osseous channels of the human calvaria appear similar to the channels previously identified in the mouse calvaria and shown to participate in immune cell trafficking between the calvarian bone and the dura mater (Cai et al., 2019; Herisson et al., 2018; Rindone et al., 2021) and in the B-cell immune surveillance of the brain (Brioschi et al., 2021; Cugurra et al., 2021).

In summary, integration of state-of-the-art LSFM and MR imaging methods provided a revised and more complete view of the vascular architecture of the dura mater. Our findings demonstrate a conserved functional anatomy of dural lymphatics between mice and humans and suggest that MLVs provide a region-specific drainage of the glymphatic outlow from the dura mater to cervical LNs. Mice can thus be reliabily used to predict the pathophysiological contribution of the dural veno-lymphatic complex and test lymphatic-targeted drugs in models of human neurological diseases.

## Material and methods

### Study approval

All in vivo procedures used in this study complied with all relevant ethical regulations for animal testing and research, in accordance with the European Community for experimental animal use guidelines (L358-86/609EEC). The study received ethical approval by the Ethical Committee of INSERM (n°2020071714182580) and the Institutional Animal Care and Use Committee of ICM.

### Animals

Male and female C57BL/6J mice, *Vegfr3YFP* (Calvo et al., 2011), or *Vegfr3-CreERT2; mTmG* (unpublished, provided by Pr. Jason Butler, Hackensack University Medical Center, NJ, USA) mice between 2 and 4 months of age were used for all experiments.

### Human patients

Procedures in humans have been approved by our institutional review board (IRB #CRM-2111-216). After informed written consent, we retrospectively collected the clinical and radiological data of six patients who underwent MR imaging with gadobutrol injection according to the protocol described below. Five of them were explored for dural venous stenoses and one for multiple sclerosis. Radiological data were then anonymized, and post-processing was performed by two experienced neuroradiologists.

### CSF Injections of tracers in mice

Intra-cisterna magna, thoracic-lumbar and lumbar-sacral level of injection was performed in adult male and female C57BL/6J and *Vegfr3YFP* mice of 8–10-week of age. Mice were injected IP with Buprecare® solution and anaesthetized by Isoflurane gas (2–3%). Mice are maintained at the head or at the vertebral level of injection with a stereotaxic apparatus (Stoelting). The skin was incised at neck level, Th12-L1 (thoracic-lumbar injection) or L6-S1 (lumbar-sacral injection) vertebrae levels. Muscles covering neck or column were moved to the side until the apparition of Dura mater. Meninges were incised using 30-gauge needle. Two or eight microliters of ovalbumin (2□mg/ml) (Ovalbumin Alexa Fluor™ 555 Conjugate (OVA-A^555^; O34782, Invitrogen) was injected through a microcapillary (Glass Capillaries; GC120-15, Harvard Apparatus) connected to a Hamilton syringe (10µl). The microcapillary was introduced into one side of the spinal cord parenchyma or above the dura mater at the cisterna magna level. To avoid the release of OVA-A^555^ during the injection, a surgical glue was added to close the incision around the glass capillary. Injections were performed slowly (1 µl/min). Once injection was finished, the capillary was maintained for 2 min before retraction and a surgical glue was added to close the hole made by the capillary, however some tracer leak always occurred despite these precautions. Tissue incisions were closed with Michel Suture Clips (7.5L×L1.75Lmm; 12040-01, Fine Science Tool). After 15, 45 or 90 min, mice were euthanized and perfused as described.

### Tissue preparation and decalcification

Mice were given a lethal dose of Sodium Pentobarbital (Euthasol Vet) and perfusion-fixed through the left ventricle with 10 ml ice-cold PBS then 20 ml 4% paraformaldehyde (PFA) in PBS. To dissect the head and the vertebrae, the skin was completely removed, all the organs were discarded, and the ribs were removed to keep only the vertebral column from the cervical part until the lumbar part with the spinal cord inside. All the surrounding tissues including muscles, eyes, salivary glandes and ligaments were maintained around the skull and the vertebral column. All samples were decalcified for 3 weeks in 10% EDTA in 4% paraformaldehyde/PBS. The head was cut with a microtome blade along coronal, horizontal or sagittal axes into either 3 pieces corresponding to the cribriform plate, cavernous sinus, jugular foramen regions, or 2 pieces corresponding to either the dorsal-versus ventral or left-versus-right halves of the head. The spine was cut along the coronal or sagittal axis into pieces of about 0.8□cm thick (2–4 vertebrae) corresponding to the cervical and the sciatic regions. The different samples segments were immediately immersed in ice-cold 4% PFA, fixed overnight at +4□°C, washed in PBS, and processed for staining.

### Paraffin section immunolabeling and imaging

Vertebrae were dehydrated through ethanol, cleared in xylene and embedded in paraffin. Serial cross sections (5□µm thick) were immunostained with rabbit anti-mouse LYVE1 (1:100) polyclonal antibody (11-034, AngioBio Co). DAB (3,3′ -Diaminobenzidine) staining was performed using the biotin avidin complex kit (PK-6100, Vectastain®Vector). Masson’s trichrome staining was carried out using the Masson Trichrome Kit (BioGnost®-Ref. MST-100T). Hematoxylin (5L) was used for counter staining. HRP-labeled paraffin sections were analyzed with a Zeiss Axio Scope.A1.

### Sample pre-treatment for iDISCO^+^ protocol

We used a clearing protocol based on methanol dehydration called the immunolabeling-enabled three-dimensional imaging of solvent-cleared organs (iDISCO^+^, http://www.idisco.info) (Renier et al., 2014). The steadily increasing methanol concentrations result in modest tissue-shrinkage (about 10%), while the “transparency” of tissues, such as the adult mouse brain, is increased. In detail, fixed samples were dehydrated progressively in methanol/PBS, 20, 40, 60, 80, and 100% for 1□h each (all steps were done with agitation). They were then incubated overnight in a solution of methanol 33%/dichloromethane 66% (DCM) (Sigma 270997-12X100□ML). After 2 □×□1□h washes with methanol 100%, samples were bleached with 5% H_2_O_2_ in methanol (1□vol 30% H_2_O_2_/5□vol methanol) at 4 °C overnight. After bleaching, samples were rehydrated in methanol for 1□h each, 80%, 60%, 40%, 20%, and PBS all steps were done with agitation). Samples were washed rapidly with PBS then incubated 2□×□1□h in PTx2 (PBS/0.2% Triton X-100). At this step they were processed for immunostaining.

### iDISCO^+^ immunolabeling protocol

Pretreated samples were incubated in 20ml glass bottle (DWK986546, Merck) PBS/0.2% Triton X-100/20% DMSO/0.3□M glycine at 37□°C for 24□h, then blocked in PBS/0.2% Triton X-100/10% DMSO/6% Donkey Serum at 37 °C for 24 h. Samples were incubated in primary antibody dilutions in PTwH (PBS/0.2% Tween-20 with 10 mg/ml heparin)/5% DMSO/3% Donkey Serum at 37 °C for 21 days. Samples were washed five times in PTwH until the next day, and then incubated in secondary antibody dilutions in PTwH/3% Donkey Serum at 37□°C for 14 days. Samples were finally washed in PTwH five times until the next day before clearing and imaging. We used the primary antibodies: Goat anti–mouse CD31 (AF3628; R&D Systems, 1:1000), Chicken anti–GFP (GFP10-20; AVES, 1:2000) Goat anti–mouse podocalyxin (AF1556; R&D Systems, 1:1000) Rabbit anti–mouse LYVE-1 (11-034, AngioBio, 1:800), Goat anti– human PROX1 (AF2727; R&D Systems;1:1000), Rabbit anti-mouse TUJ1 (802001, Biolegend; 1:2000), Rabbit anti–human vWF (A0082, Agilent, 1:300). Primary antibodies were detected with the corresponding Alexa Fluor -555, -647, -790 conjugated secondary antibodies from Jackson ImmunoResearch at 1/1000 dilution.

### iDISCO^+^-tissue clearing

After immunolabeling, samples were dehydrated progressively in methanol in PBS, 20, 40, 60, 80, and 100% each for 1□h (all steps were done with agitation). They were then incubated overnight in a solution of methanol 33%/DCM 66% followed by incubation in 100% DCM for 2□×□1hr to wash the methanol. Finally, samples were incubated in dibenzyl ether (DBE) (without shaking) until cleared (overnight) and then stored in DBE at room temperature before imaging.

### LSFM and stereomicroscope imaging

Cleared samples were imaged in transverse orientation with a LSFM (Ultramicroscope II, LaVision Biotec) equipped with a sCMOS camera (Andor Neo) and a 4□×□/0.3 objective lens (LVMI-Fluor 4□×□/0.3 WD6, LaVision Biotec). Version v144 of the Imspector Microscope controller software was used. The microscope chamber was filled with DBE. We used single sided 3-sheet illumination configuration, with fixed x position (no dynamic focusing). The light sheet was generated by LED lasers (OBIS) tuned to 561Lnm 100□mW and 639Lnm 70□mW (LVBT Laser module 2nd generation). The light-sheet numerical aperture was set to 0.03. We used the following emission filters: 595/40 for Alexa Fluor-568 or -555, -680/30 for Alexa Fluor-647 and 830/780 for Alexa Fluor-790. Stacks were acquired using 4.5 μm z steps and a 30 ms exposure time per step, with a Andor CMOS sNEO camera. The ×2 optical zoom was used for an effective magnification of (×8), 0.8□µm/pixel. Mosaic acquisitions were done with a 10% overlap on the full frame.

Fluorescent stereo micrographs were obtained with AxioZoom.V16 fluorescence stereo zoom microscope (Carl Zeiss) equipped with an ORCA-Flash 4.0 digital sCMOS camera (Hamamatsu Photonics) or an OptiMOS sCMOS camera (QImaging).

### LSFM-image processing and analysis

For display purposes, a gamma correction of 1.47 was applied on the raw data obtained from the light-sheet fluorescent microscope.

Images acquired with Imspector acquisition software in tif fomat was converted with Imaris File Converter to IMS files. Mosaics were reconstructed with Imaris stitcher; then Imaris software (Bitplane, http://www.bitplane.com/imaris/imaris) was used to generate the orthogonal projections of data shown in all figures, perform area segmentation on a stack of image slices and produce videos.

### MR Imaging protocol

The following sequences were performed using MRI 3.0 Tesla Magnetom Skyra (Siemens Healthineers). The following sequences were performed in all patients after Gadobutrol (0.1 mmol/kg body weight, i.v., Bayer Health Care, NDC 50419-325-12) injection: Contrast-enhanced MR venography with elliptical-centric technique (3D sagittal acquisition, field-of-view 250 mm^2^, 208 contiguous 0.70 mm slices, TR/TE = 3.57/1.36 msec, acquisition time 6 min); Coronal Whole Brain 3D FLAIR (Coronal 3D acquisition, field-of-view 256 mm^2^, 192 contiguous 1 mm slices, TR/TE = 5000/375 msec, acquisition time 6 min); Whole Brain T1 SPACE DANTE acquisition (Sagittal 3D acquisition, field-of-view 256 mm^2^, 224 contiguous 0.80 mm slices, TR/TE = 7000/22 msec, acquisition time 6 min).

### MRI post processing

3D-Slicer platform was then used for semi-automatized signal intensity-based thresholding and segmentation of the native sequences. The whole Brain T1 SPACE DANTE sequence was post-processed and two manual segmentations were performed to extract the dural lymphatics and the parasinus enhancement at the dural sinuses and the cLNs, and the internal carotid arteries (iCA). In one patient with multiple sclerosis, a 3D TOF (Time of Flight) sequence before Gadobutrol injection was also performed and was used to reconstruct the arterial system. 3-dimensional reconstruction of the dLVs and of their related cervical lymph nodes and of the ICA were then obtained. The contrast-enhanced MR venography with elliptical-centric technique was than post-processed and merged with the 3D-reconstruction of the dLVs to confirm that lymphatic vessels were not misdiagnosed as small slow flow veins and to detail the veno-lymphatic relationships in humans. The combination of these two-reconstruction allowed a 3D visualization of the whole vascular system in Humans.

### Flow-cytometry analysis of LN immune cells

At 90min after spinal intrathecal injection of OVA-A^488^, mice were anesthetized with Ketamine/Xylazine. Lymph nodes (mandibular, accessory mandibular, deep cervical, sciatic, lumbar lymph nodes) were dissected and processed as previously described (Geraldo et al., 2021). LNs were digested with DMEM containing 2.5 mg/ml collagenase D, and 5 U/ml DNase I for 20 min at 37°C. The digested tissue was passed through a 40μm nylon cell strainer (Falcon) and red blood cells were lysed (Red Blood Cells Lysis buffer, Merck). After blocking with mouse FcR Blocking Reagent (MACS Miltenyi Biotec), single cell suspensions were incubated with anti-CD45 BUV805 (Clone 30-F11, BD), anti-CD11b BV421 (Clone M1/70, BD), anti-CD11c APC (Clone N418, BD) antibodies. As a control, cells were stained with the appropriate isotype control. Data acquisition was performed on BD LSRFortessa X20 and analysis was performed with FlowJo_V10.

### Statistics

No statistical methods were used to predetermine sample size. Five to six mice were analyzed by experimental group (*n*L=L5–6 mice/group). The investigators were blinded during experiments and outcome assessment.

Statistical data analysis was performed with the Prism 6.0 software (GraphPad). For discrete variables (immune-cell %), data are presented as mean standard error of the mean (SEM). A two-tailed, unpaired Mann–Whitney U test was done to determine statistical significance between two groups. For comparison between more than two groups, the one-way ANOVA test was performed, followed by Turkey’s multiple comparison test. Differences were considered statistically significant if the *p* value was <0.05 (\**p*□<□0.05, \*\**p*□<□0.01).

### Graphic design

3D-Schemas of the mouse head vasculature were generated using 3D-Blender, a free and open-source 3D computer graphics software toolset, from horizontal, sagittal and coronal representations of the rat dural venous anatomy (Scremin, 2004) and from horizontal, sagittal and coronal head sections labeled with vascular markers.

## Online supplemental material

**Videos 1-6.** LSFM coronal (1, 3, 4, 5) and sagittal (2, 6) images of clarified half-heads performed 45 min after intrathecal injection of CSF tracer (OVA-A^555^) into the caudal spine, with additional immunolabeling with anti-LYVE1 antibodies in (4-6).

**Video 1.** OVA-A^555^ tracer (white) labels phagocytic cells located along perivascular spaces of the brain vasculature as well as along the superior sagittal sinus in the dura mater.

**Video 2.** OVA-A^555^ tracer (white) labels brain meninges and, outside of the skull, the nasal cavity, the nasopharynx and the cervical LNs.

**Video 3.** Superficial cervical LNs are labeled by OVA-A^555^ tracer (white).

**Video 4**. LYVE1^+^ MLVs (green) are localized at the carotid foramen and are associated with OVA-A^555^ deposits around the cavernous sinus.

**Video 5.** At the entry point of the ophthalmic emissary vein into the anterior cavernous sinus, LYVE1^+^ lymphatic capillaries (green) are in contact with the OVA-A^555^ labeled-wall of the cavernous sinus (magenta). OVA-A^555^-positive phagocytic cells are visible inside the dural lymphatic capillary.

**Video 6.** LYVE1^+^ MLVs are localized around the olfactory bulbs, but no LYVE1+ lymphatic connection is observed through cribriform foramina with the nasal cavity. A dense network of OVA-A^555^-positive vascular like structures is observed in the nasal cavity.

## Acknowledgements

This work was supported by ANR-17-CE14-0005 BrainWash, ANR-20-CE16-0027-01 Lymbrain, BBT.1300 Venolymphatic, and Programme Capes Cofecub 2021 projet n° 44999PG to JLT). The authors are funded by the following sources: LJ (BrainWash), JB (Capes Cofecub), SLenck (BBT.1300 Venolymphatic; INSERM CIHU-2021), LG (European Society of Cardiology (ESC) Basic Research Fellowship), MREK (Lymbrain), SLehericy (Investissements d’Avenir, IAIHU-06 (Paris Institute of Neurosciences – IHU)), ANR-11-INBS-0006), and AE (European Research Council (ERC) grant agreement No. 834161). All animal work was conducted at the PHENO-ICMice facility. The Core is supported by 2 ‘Investissements d’Avenir’ (ANR-10-IAIHU-06 and ANR-11- INBS-0011-NeurATRIS) and the Fondation pour la Recherche Médicale. We are greatly indebted to the ICM-Quant and -Histomics platforms and Paris Cardiovascular Research Center (PARCC) Flow and Image Cytometry facility (Dr. Camille Knosp) for their technical assistance. We are grateful to: Phil Colsh (Yale Neurology) for his advice and careful reading and editing of the manuscript; Rejane Rua (Centre d’Immunologie de Marseille Luminy) for her critical reading of the manuscript; Alexis Vaussy (Siemens, Healthineers) and Siemens company for their technical collaboration to improve the 3D T1 SPACE DANTE sequence.

## Author Contributions

LSFM data (LJ, JB), VWI data (S Lenck, S Lehericy, FB, CC), LJ, IHC on sections (JB, MREK), flow cytometry (LJ, LG, YX), iconography (JMT), resources (SH, MCP, AE and JLT), manuscript writing (LJ, JBN, SL, AE and JLT). All authors read and edited the manuscript.

## Competing interests

The authors declare that they have no conflict of interest.

## Supplementary data

**Figure S1. Bidirectional propagation of OVA-A^555^ tracer after spinal intrathecal injection.**

**A, B** Schema of the tracer distribution after Th-Lb or Lb-Sa injection. Mice were sacrificed at 15 or 45 min after tracer injection. Macroscopic imaging of OVA-A^555^ (white)/ FITC-dextran (green) demonstrated a similar pattern at different levels of the caudo-rostral CNS axis (white) and a bi-directional propagation of tracers along the spinal cord (white arrows). SC: spinal cord.

**C-F** Cross sections of the cervical and sacral spine showing ink (black) propagation in the pia mater along the rostral-caudal axis. Samples were imaged immediately after dissection (**C**, **E**) and following paraffin embedding and sections of the cervical and sacral spine (ink **D, F**).

**G** Cross section of the posterior head showing ink deposits lining the brainstem and inside the dcLNs (green arrowheads). BS: brainstem.

**H** LSFM coronal view of the CSF tracer pattern (OVA-A^555,^ white) in the brain pia mater, in the perivascular space and the scLNs (double green arrowheads). NP: nasopharynx. **i** Sagittal section of the forehead showing ink deposits in the pia mater and throughout the outflow pathway of the olfactory epithelium. Cx: cortex, OB: olfactory bulb, OE: olfactory epithelium. A: anterior, D: dorsal, L: lateral, P: posterior, Th-Lb: Thoraco-lumbar, V: ventral. Scale bar: 500 μm (**A**-**F**); 2 mm (**G**-**H**).

**Figure S2. OVA-A^555^ tracer accumulation around dural sinuses and cerebral veins.**

**A, B** Coronal (**A**) and sagittal (**B**) LSFM views of the peri-sinus distribution of tracer deposits (blue arrowheads) along the transversal sinus (vWF+, blue, blue arrowhead).

**C, D** Caudal (**C**) and rostral (**D**) convergence of sinuses show MLVs (green) associated with tracer deposits (magenta).

**E.** Lateral view of the distribution of tracer (magenta) in the peri-sinus area of the transverse, sigmoid and petrosquamous sinuses (blue). TS: transverse sinus, SS: sigmoid sinus, PSS: petrosquamous sinus.

**F** Blood vessels of the jugular foramen region (CD31^+^/PDXL^+^, red). A: anterior, COS: caudal confluence of sinuses; Cx: cortex, D: dorsal, L: lateral, P: posterior, jugv: jugular vein. Scale bar: 200 μm (**A**-**D**); 500 μm (**E**, **F**).

**Figure S3. Lymphatic drainage in the caudal region of cavernous sinuses.**

**A** Schematic lateral view of the meningeal vasculature and position of lymphatic uptake hotspots (1-3) in the middle fossa of the skull base. cav: cavernous sinus, ic: internal carotid artery, ipets: inferior petrosal sinus, iptgev: interpterygoid emmissary vein, spets: superior petrosal sinus.

**B** Blood vessels (CD31^+^/PDLX^+^, red) around the pituitary gland (PG) and the cavernous sinus (cav, light-blue dashed line).

**C** Horizontal view of tracer deposits (magenta) and vWF^+^ dural sinuses and veins (blue) showing localization of lymphatic uptake hotspots (1-3). CC: carotid canal, icav: inter-cavernous sinus, ipets: inferior petrosal sinus.

**D-F** Vascular afferences to the cavernous sinuses. Blood vessels (CD31^+^/PDLX^+^, orange in **D**, **E;** vWF^+^, blue in **F**). Coronal views of the transverse sinus connection with the inter-cavernous sinus (**D**) and of the posterior basal vein (pbv) with the cavernous sinus (**E**). Sagittal view showing the connection of inferior petrosal sinus and basilar vein with the cavernous sinus (**F**).

**G-I** Carotid canal area. Blood vessels (CD31^+^/PDLX^+^, orange in **G**) and lymphatics (LYVE1^+^, green in **H, I**) on coronal (**G**, **I**) and sagittal (**H**) views of the carotid canal. **I**: coronal section in **H**. Tracer deposits in the pia and along the carotid artery (yellow in **G**). Lymphatic vessels (green) around the carotid canal in **H**, **I**. Scale bar: 500 μm (**B**-**G**, **I**); 1500 μm (**H**).

**Figure S4. Lymphatic drainage in the rostral region of cavernous sinuses.**

**A** Ventral view of cerebral veins and dural sinuses (vWF^+^, blue) and peri-sinus tracer deposits (magenta). Tracer accumulates at confluence points of the cavernous sinus with the ophthalmic and olfactory emissary veins (4, 5) and with the inferior olfactory sinus (6). The middle meningeal vein and the pterygoid venous plexus converged with the ophthalmic emissary vein at the anterior lacerated foramen (dotted line). Anterior lacerated foramen: alf, middle meningeal vein: mmv, of: optic foramen, ON: optic nerve, ophev: ophthalmic emissary vein, ptglp: pterygoid venous plexous, rrhv: rostro-rhinal vein, *V*: cranial nerve 5 (Trigeminal).

**B-D** Ophthalmic emissary vein region (4). Cranial nerves (**B**), blood vessels (**C**) and lymphatics (**D**). Coronal views of the pattern of cranial nerves (TUJ1^+^, yellow), blood vessels (CD31^+^/PDXL^+^, orange) and lymphatics (LYVE1^+^, green). Cranial nerves were devoid of both tracer deposits (**B**). The optic nerve showed a perivascular network (**C**). Lymphatics located in the dural cavernous sinus connect with the lymphatic network of the nasopharynx (white arrow in **D**). ON: optic nerve, *III*: cranial nerve 3 (Oculomotor), *IV*: cranial nerve 4 (Trochlear), *V*: cranial nerve 5 (Trigeminal), *VI*: cranial nerve 6 (Abducens).

**E-G** Olfactory emissary veins region (4, 5). Blood vessels (**E**) and lymphatics (**F, G**). Coronal views of the pattern of blood vessels (CD31^+^/PDXL^+^, orange) and lymphatics (LYVE1^+^, green) in the orbital cavity and the cavernous sinus. Tracer deposits (magenta) concentrate in the rostral part of the cavernous sinus and the caudal part of the orbital cavity. Lymphatics from the orbital cavity run along the olfactory and ophthalmic emissary veins and enter the skull through the anterior lacerated foramen to contact the peri-sinus of the dural cavernous sinus (4, 5).

**H-K** Inferior olfactory sinus (IOS) region (6). Cranial nerves (**H**), blood vessels (**J**) and lymphatics (**J, K**). Coronal (**H-J**) and sagittal (**K**) views of the pattern of nerves (TuJ1^+^, yellow), blood vessels (vWF^+^, blue) and lymphatics (LYVE1^+^, green) in the most rostral part of the cavernous sinus. Tracer deposited along the ON until the retina, and in the peri-sinus of cavernous sinus (**H**). IOS runs at the surface of the olfactory bulb (OB) to reach the superior olfactory sinus (SOS) in the calvaria dura (**I**). Lymphatic tracer uptake was detected in the peris-sinus of the cavernous sinus (**J, K**). Scale bar: 500 μm (**A**-**K**).

**Figure S5. CSF drainage through the cribriform plate and inside the nasal cavity. A**

Lymphatics in the cribriform plate. Tracer deposits at the cribriform plate (CP) border and in PROX1^+^ lymphatics around olfactory nerve foramina (orange arrowheads).

**B-E** Lymphatics in the nasal cavity. Sagittal (**B, C**), and coronal (**D, E**) views showing tracer deposits (magenta in **B**, **C**, **E**; ink in **D**) and pattern of PROX1 (**B**, green), VEGFR3-GFP (**C**) and LYVE1 (**D**, **E**) lymphatics. **B**, **C** PROX1^+^ lymphatic vessels in the turbinate, on the outer side of the cribriform plate (white arrows in **B**), and VEGFR3^+^ (green) vessels transporting OVA-A^555^ tracer in the olfactory epithelium (yellow arrowheads in **C**). **D** Contiguous paraffin sections of the nasal cavity showing ink tracer is drained by narrow LYVE1^-^ vessels and collecting LYVE1^+^ lymphatics in the respiratory epithelium. **E** LYVE1^+^ lymphatics, including tracer-draining vessels (white, arrowheads) around the nasopharynx.

**F** Lymphatic drainage from the orbital cavity. Coronal view showing tracer deposits (magenta) and pattern of LYVE1^+^ lymphatics (green). Lymphatic-tracer uptake is detected around the nasal lacrimal sac (white, arrows).

**G** Synthetic schema of lymphatic circuits of CSF drainage in the mouse head. Cerebral veins and dural sinuses (deep blue) drain blood from the following brain regions: dorsal (orange), anterior (pink), basal (green) and caudal (light blue). Their associated perivenous glymphatic efflux provides a brain region specific-information to the peri-sinus and lymphatic components of the dura mater. Lymphatic uptake hotspots are annotated with numbers from caudal to rostral (1-6). Light green: MLVs; bluish green: nasal cavity lymphatics; deep green: extracranial collecting lymphatics and LNs; light blue: efferent veins; red: internal carotid artery; salmon: nasopharynx. amLN: accessory mandibular lymph node, cav: cavernous -sinus, cpf, cribriform plate foramina, dcLN: deep cervical lymph node, jugv: jugular vein, pfv: posterior facial vein, ic: internal carotide, IOS: inferior olfactory sinus, IPETS: inferior petrosal sinus, jf: jugular foramen, OE: olfactory epithelium, psf: petrosquamous fissure, PSS: petrosquamous sinus, mLN: mandibular lymph node NP: nasopharynx, RCS: rostral confluence of sinuses, RE: respiratory epithelium, SOS: superior olfactory sinus, SS: sigmoid sinus SSS; superior sagittal sinus

Scale bar: 300 μm (**A**); 150 μm (**B**-**F**).

**Figure S6. Sacral spinal cord outflow.**

**A, B** Macroscope imaging of the sacrococcygeal region after i.c.m injection of OVA-A^555^ tracer. **A** Tracer deposits were detected at inter-vertebral spaces (white arrows, S3-S4, S4-Co1, Co1-Co2) and in the sciatic LNs (siLN, green arrowhead). **B** In the peritoneal cavity, the tracer was detected in collecting lumbar LNs (lbLN) and renal LNs (reLN) (white, green arrowhead). asterisk: vertebral body.

**C-E** LSFM sagittal (**C**) and coronal (**D, E**) views of clarified sacral vertebrae, spinal cord and dura mater. **C** Outflow of tracer (red) in the epidural space (white arrows) between S3-S4 and S4-Co1. asterisk: vertebral body. **D**, **E** Pattern of LYVE1^+^ lymphatics (green) and tracer (red) in the S3-S4 region. Tracer deposits (red) were detected in the vertebral canal and in siLNs (green arrowheads in **D**). Tracer accumulated in the epidural space, including in LYVE1^+^ lymphatics (white, arrow in **E** (magnification of dotted frame in **D**). A: anterior, D: dorsal, L: lateral, V: ventral, S: sacral vertebrae.

**F** Graphics representing the percentage of tracer-labeled macrophages (CD45^+^/CD11b^+^/CD11c^-^) in the siLNs and lbLNs of control mice after spinal OVA-A^555^ tracer injection.

**G-K** Light field microscope images of paraffin cross sections of S3-S4 vertebrae from mice administered with intraspinal injection of Indian ink tracer. **G** Large-field image with localization of highly magnified areas shown in **H-K**. Tracer was detected at the pial surface of the spine and the ependymal border (**H**), as well as in phagocytic cells in the subarachnoid space (**I, J**) and the epidural space (**K**). Note the thin monolayer of dura mater (black arrow in **I**) and the accumulation of tracer-containing mononuclear cells in sub-dural arachnoid sacs (black arrow in **J**). Scale bar: 2 mm (**A**, **B**); 500 μm (**C**-**G**); 10 μm (**H**-**K**).

**Figure S7. Meningeal and skull vascular MR-Imaging in humans.**

**A** Trans-calvarian connections at the superior longitudinal sinus (SLS). A few trans-calvarian veins (TcV) connecting the SLS with subcutaneous veins (SuV) and the systemic circulation were associated with tiny non-venous channels filled with parasinus fluids (red asterisk).

**B** Native 3D T1 SPACE DANTE sequence after gadobutrol injection in a coronal plane crossing the cribriform plate (CP) and the maxillary sinuses (MS). Gadobutrol enhancement was detected intracranially along the SLS (white arrow) and the perivenous space of several cortical veins (white arrowhead). Extracranially, the olfactive epithelium was strongly enhanced after gadobutrol injection (OE), but no gadobutrol signal was observed from the dura mater across the CP (light blue arrow). Scale bar: 1 cm (**A**, **B**).

**C** Schematic representation of the dorso-basal and anterior veno-lymphatic complexes focusing on the lymphatic outflows. Gadobutrol parasinus enhancement was depicted around most dural sinuses, including the SLS, the straight sinus (StS), the transverse sinus (TS) and the sigmoid sinuses (SS) and the cavernous sinus (CS). Significant gadobutrol flow was detected in the carotid canal along the internal carotid artery (ICA) or follows trans-foraminal routes along the trigeminal nerve branches (grey) through the superior orbital fissure (SOF), the foramen rotundum (FR) and the foramen ovale (FO). We propose a model where, according to the region-specific venous circulation in brain tissues (orange, pink, green and light blue brain regions), perivenous glymphatic efflux provide a brain region specific-information to the peri-sinus and lymphatic components of the dura mater, thereby conferring a brain region specific immune surveillance of the brain tissues from the meninges.

## Notes

### Competing Interest Statement

The authors have declared no competing interest.

