## Supplementary figures and images for "3D-imaging reveals conserved cerebrospinal fluid drainage via meningeal lymphatic vasculature in mice and humans"

### Supplemental Fig1-Fig7

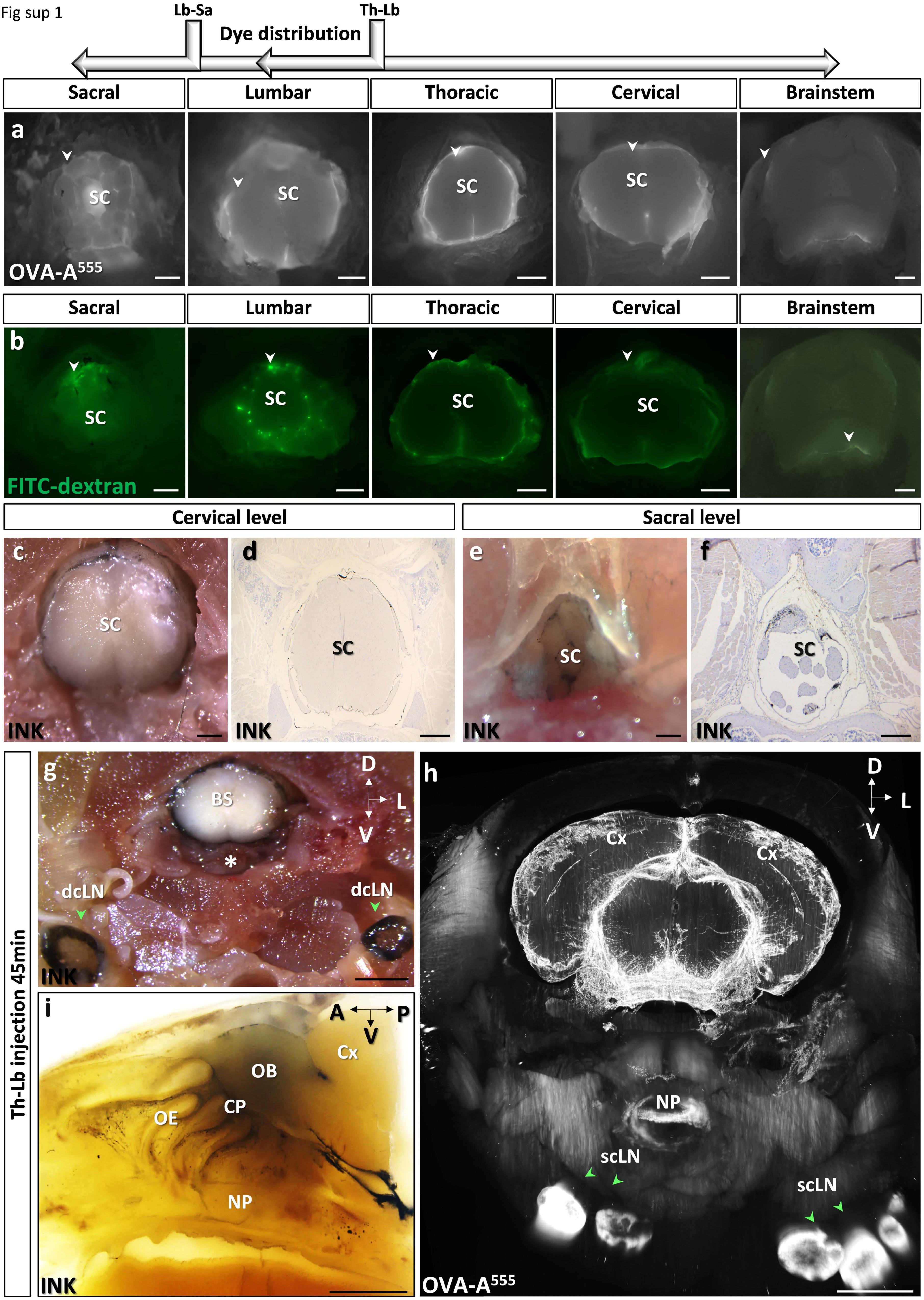

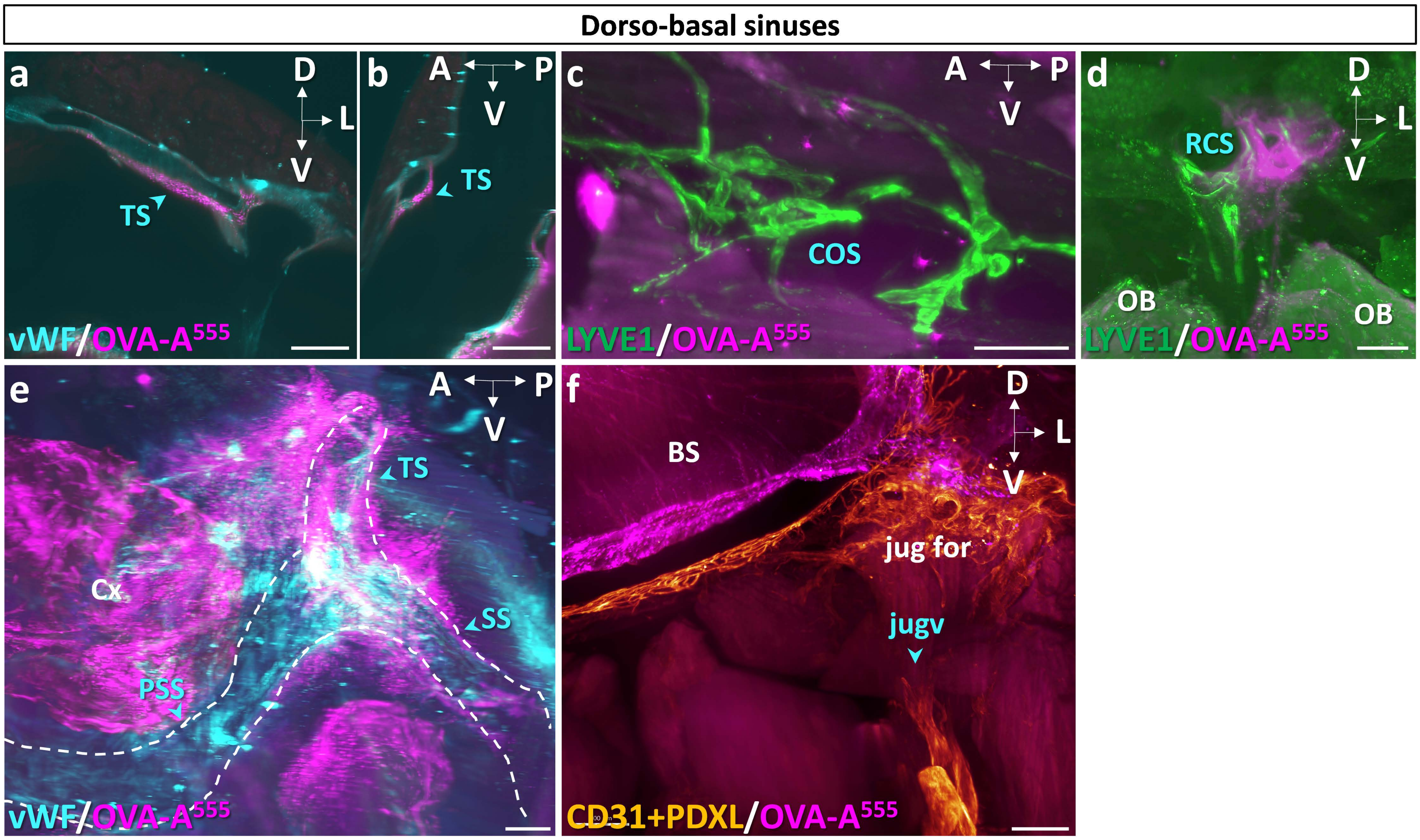

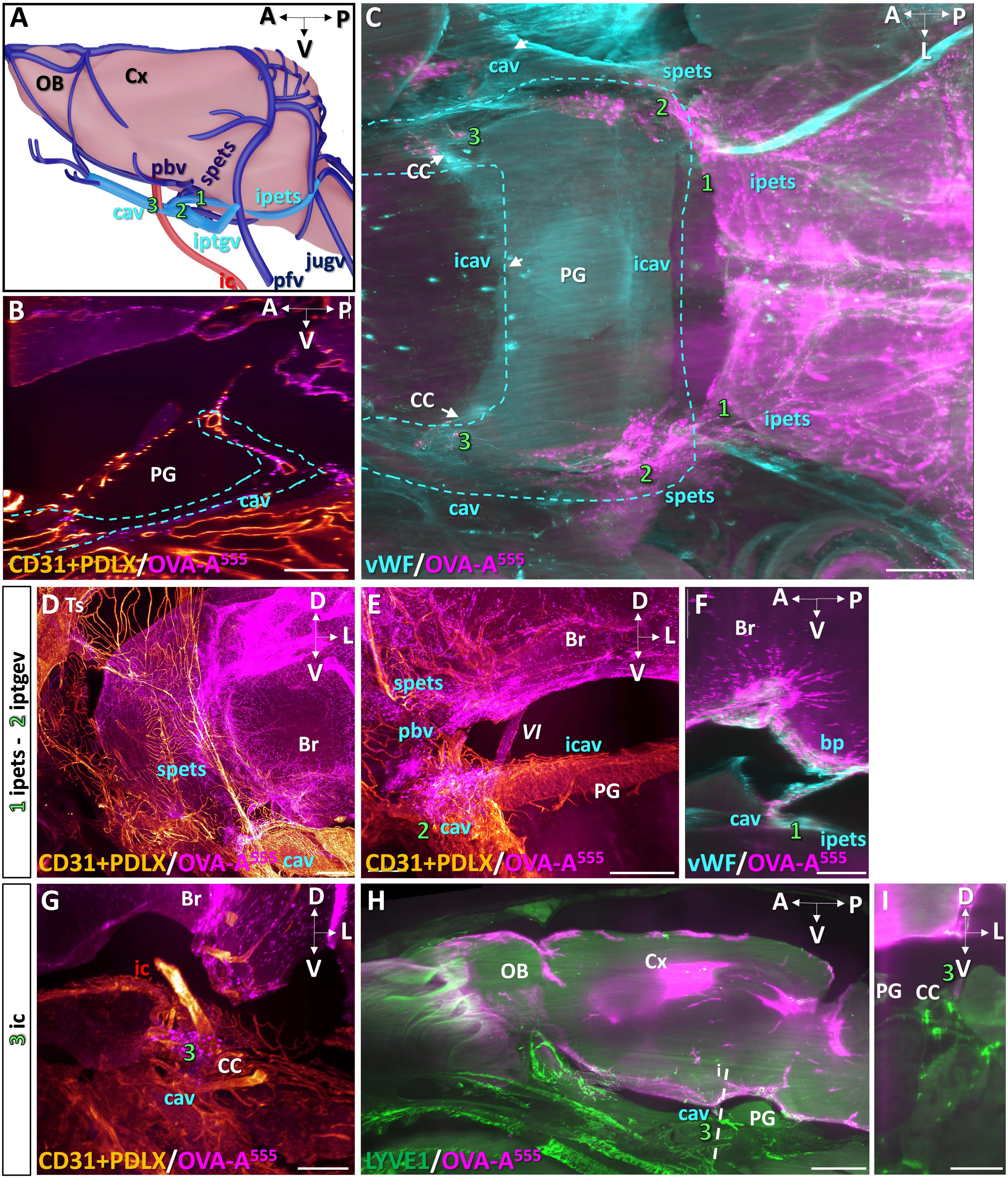

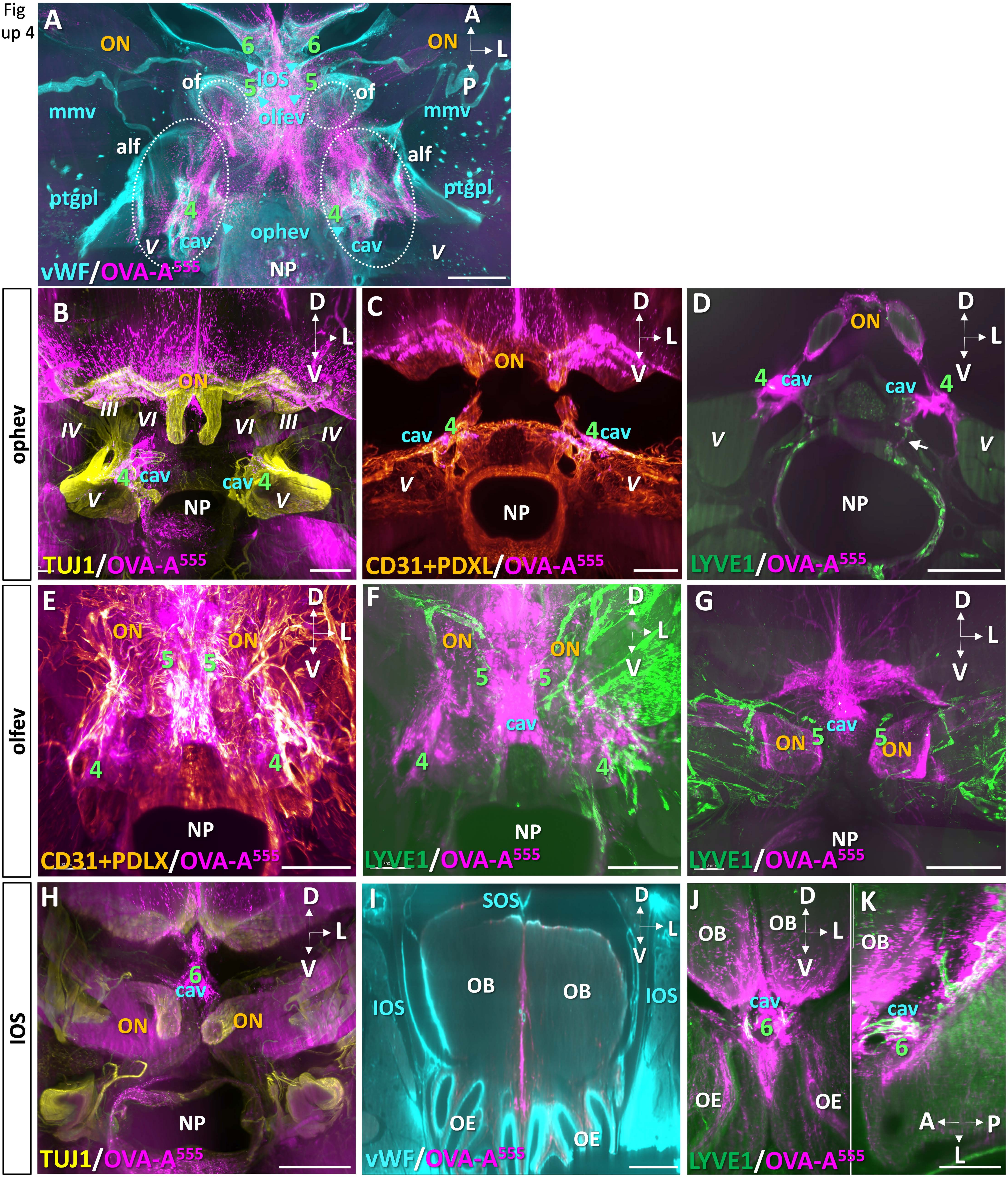

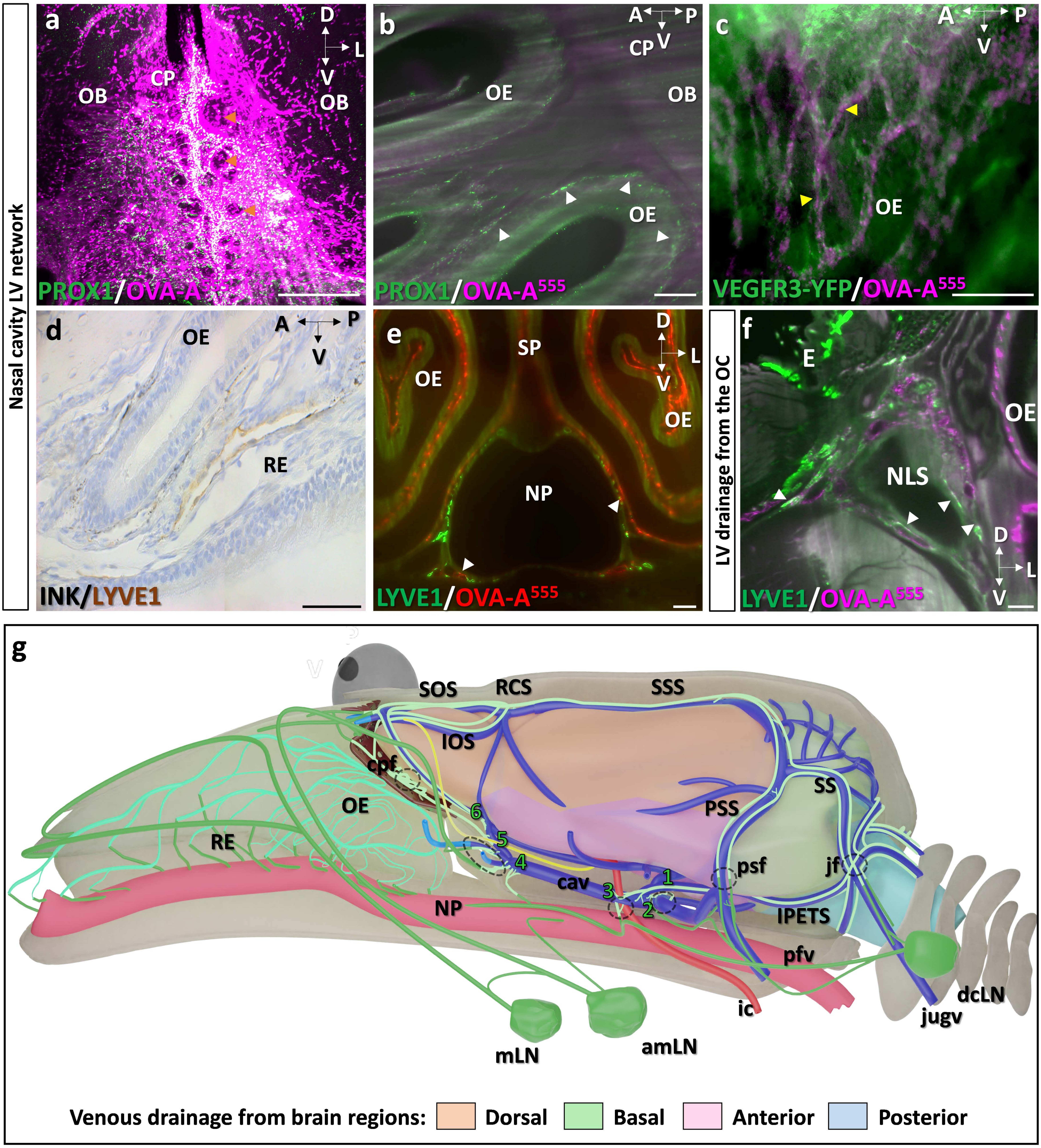

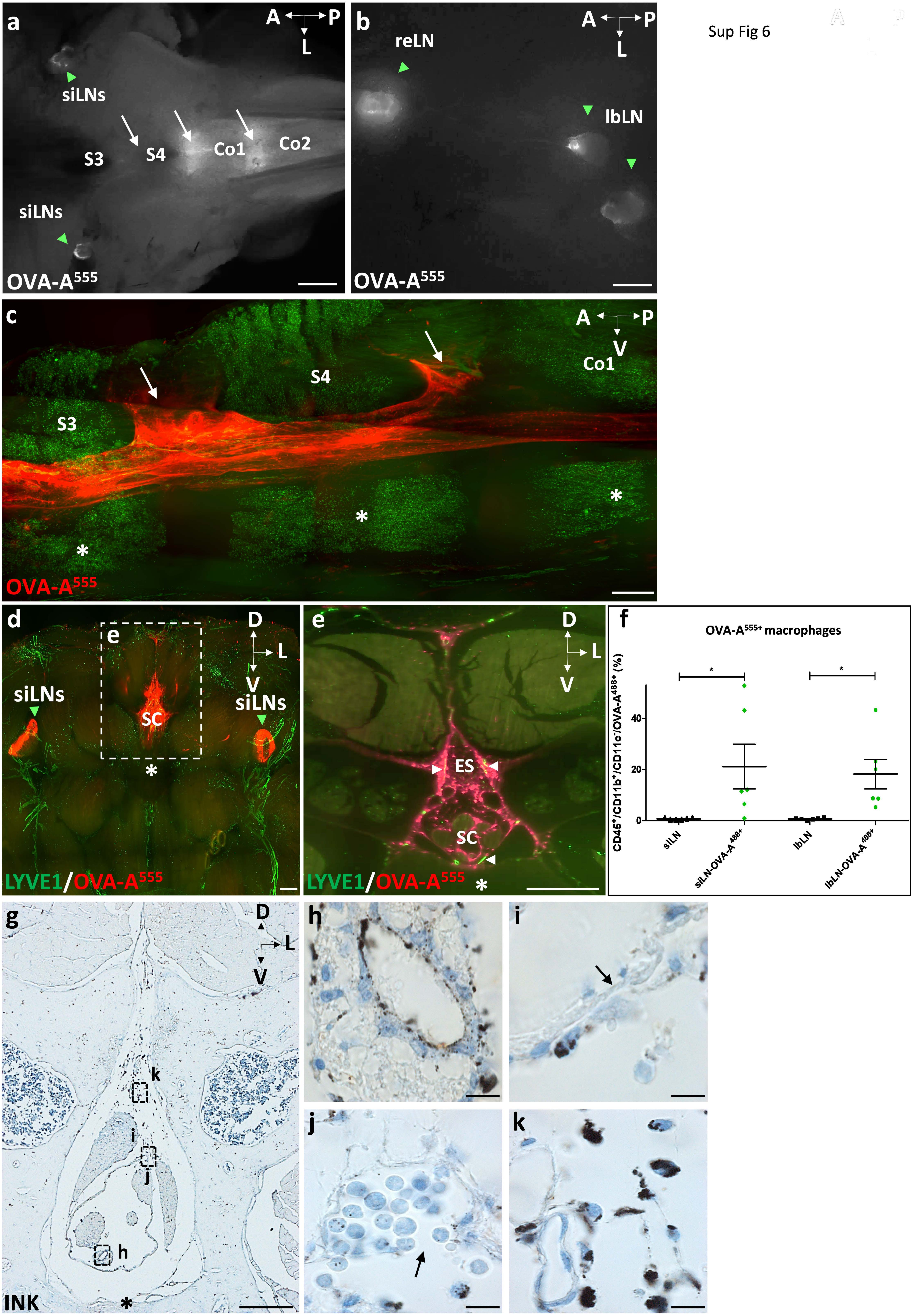

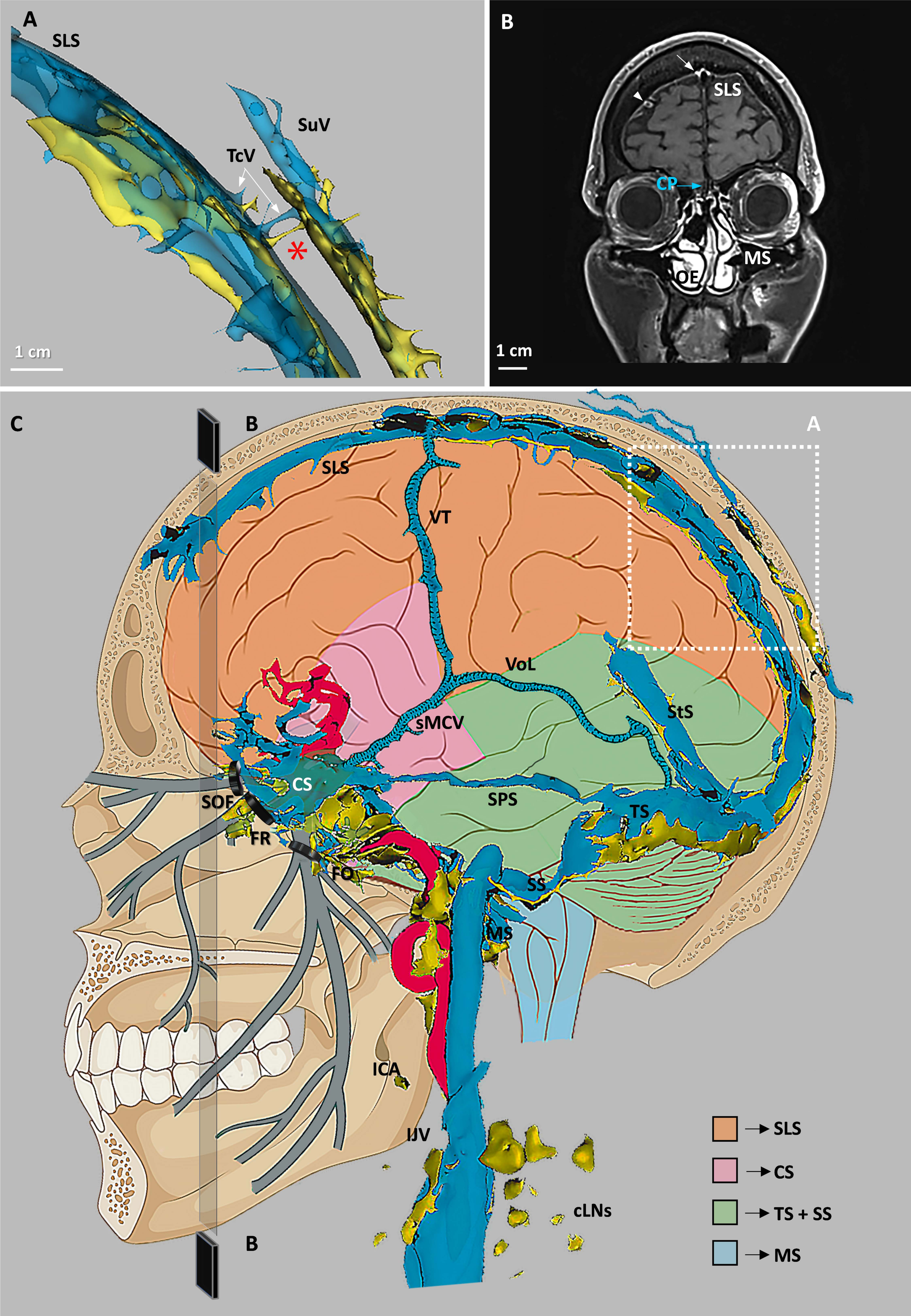
